# The transcriptional response to low temperature is weakly conserved across the *Enterobacteriaceae*

**DOI:** 10.1101/2024.06.10.598259

**Authors:** Johnson Hoang, Daniel M Stoebel

**Affiliations:** Department of Biology, Harvey Mudd College, Claremont, California, USA

## Abstract

Bacteria respond to changes in their external environment, such as temperature, by changing the transcription of their genes. We know little about how these regulatory patterns evolve. We used RNA-seq to study the transcriptional response to a shift from 37°C to 15°C in wild-type *Escherichia coli, Salmonella enterica, Citrobacter rodentium, Enterobacter cloacae, Klebsiella pneumoniae,* and *Serratia marcescens*, as well as Δ*rpoS* strains of *E. coli* and *S. enterica.* We found that these species change the transcription of between 626 and 1057 genes in response to the temperature shift, but there were only 16 differentially expressed genes in common among the six species. Species-specific transcriptional patterns of shared genes were a prominent cause of this lack of conservation. GO enrichment of regulated genes suggested many species-specific phenotypic responses to temperature changes, but enriched terms associated with iron metabolism, central metabolism, and biofilm formation were implicated in at least half of the species. The alternative sigma factor RpoS regulated about 200 genes between 37°C and 15°C in both *E. coli* and *S. enterica*, with only 83 genes in common between the two species. Overall, there was limited conservation of the response to low temperature generally, or the RpoS-regulated part of the response specifically. This study suggests that species-specific patterns of transcription of shared genes, rather than horizontal acquisition of unique genes, are the major reason for the lack of conservation of the transcriptomic response to low temperature.

**Importance:** We studied how different species of bacteria from the same Family (Enterobacteriacae) change the expression of their genes in response to a decrease in temperature. Using de novo-generated parallel RNA-seq datasets, we found that the six species in this study change the level of expression of many of their genes in response to a shift from human body temperature (37°C) to a temperature that might be found out of doors (15°C). Surprisingly, there were very few genes that change expression in all six species. This was due in part to differences in gene content, and in part due to shared genes with distinct expression profiles between the species. This study is important to the field because it illustrates that closely related species can share many genes but not use those genes in the same way in response to the same environmental change.

## Introduction

Bacteria change the levels of transcription of their genes to respond to changes in their environments. These patterns of transcription, and the role of regulatory proteins and other factors in determining these patterns have been measured thoroughly in a handful of model species, like *Escherichia coli*. The degree to which our understanding of *E. coli* gene regulation can inform our understanding of related species depends on how similar those species are. We know that the genomes of bacterial species often differ dramatically in content due to horizontal gene transfer (HGT), so it is possible that HGT may drive dramatic differences between related species in what genes are differentially expressed (DE) in response to an environmental change. Differences in transcriptome patterns between related species can also be due to shared genes that have different patterns of regulation among species. In general, we have little information about the relative roles of species-specific genes vs. species-specific regulation of shared genes in shaping how regulons differ between related species.

Changes in temperature are one challenge faced by many bacteria (1). Low temperature can affect bacteria by rigidifying the lipid bilayer, by altering the folding of proteins and RNAs, and by influencing the assembly and activity of essential macromolecular machinery like ribosomes (1). Low temperature may also serve as a cue that bacteria have moved from one environment (such as a mammalian host) into another environment (such as a surface at ambient temperature) (2).

A great deal is known about how *E. coli* responds to low temperature (1, 3). Several global regulatory factors have been implicated, including cold shock proteins (4, 5), RNA degradation enzymes like RNase R and PNPase (5, 6), nucleoid associated proteins like H-NS (7), global supercoiling levels (8), and the alternative sigma factor RpoS (2, 9). Shifts between optimal temperature (37°C) and a lower temperature have substantial impacts on the transcriptome. In a series of experiments examining shifts between 37°C and 23°C (and vice versa), White-Ziegler *et al.* documented that 423 genes, or about 10% of the *Escherichia coli* K-12 genome, are DE (2, 10). Further, they identified RpoS and H-NS as two global regulators of these transcriptional patterns (2, 11).

How relatives of *E. coli* respond to changes in temperature is much less well documented. The *Enterobacteriaceae*are a diverse group of bacteria (12) that exist in many different thermal environments (13). This group includes genera that can be found in endothermic hosts either causing an infection or in the gastrointestinal tract of healthy hosts (14). Many of these species can also be isolated from ectothermic animals, plants, soil, water, and plants (15–19). The presence of these organisms outside of endothermic hosts suggests that they naturally experience broad changes in temperature.

In general, we have little understanding of how regulatory responses evolve in bacteria and at what rate these regulons evolve. In addition to questions of rate, regulons could evolve because different species lose different genes or gain different genes via horizontal gene transfer (HGT). Regulons could also evolve because shared genes evolve different regulatory controls in different species. We do not understand the relative importance of these two modes of regulatory evolution. Investigating the tempo and mode of regulatory evolution in the *Enterobacteriaceae*will both allow fundamental understanding of bacterial evolution and allow researchers to better understand the degree to which *E. coli* serves as an appropriate model system for related bacteria.

Single species studies of the transcriptomic response to low temperature exist for several species of *Enterobacteriaceae*, including *E. coli* (2, 10), *Salmonella enterica* (20), *Klebsiella pneumoniae*(21) and *Enterobacter sp.*(22). Unfortunately, it is difficult to directly compare existing studies of individual species because of the large differences in culture conditions, overall experimental approach (e.g. microarrays versus RNA-seq), experimental details (e.g. methods of library construction for RNA-seq), and bioinformatic and statistical analyses in these studies. To directly compare the transcriptional response, the experimental and analytical approaches should be the same. In addition, batch effects in both bacterial growth and RNA-seq steps can be avoided by performing all the bacterial growth, RNA processing, library construction, and sequencing in parallel with a balanced experimental design.

In this study, we studied how five species of bacteria in the *Enterobacteriaceae*plus one closely related species responded to the shift from 37°C to 15°C, as might occur when bacteria leave a mammalian host and find themselves in a temperate external environment. We used comparative genomic tools to identify homologous genes in each species, and then used RNA-seq to examine the patterns of transcription in each species across this temperature transition. In addition, we studied the role of the alternative sigma factor RpoS in this transition in *E. coli* and *S. enterica.* We performed these studies in parallel for all strains, avoiding possible confounding differences due to experimental or analytical approaches or due to batch effects.

## Results

### Most genes are either shared across all species or are species-specific

This study worked with strains of five species of *Enterobacteriaceae*, (*E. coli* K-12*, S. enterica* serovar Typhimurium 14028s*, Citrobacter rodentium*DBS100*, Enterobacter cloacae* ATCC13047*, K. pneumoniae* MKP103) and one species in the *Yersiniaceae* (*Serratia marcescens* Db11) (Figure 1A) (12). Their genomes contained between 3968 and 4709 protein coding sequences (Figure 1B), which were the basis of all the analyses in this paper. We used xenoGI(23) to understand the evolutionary histories of the six genomes. xenoGI uses both synteny and sequence similarity to define groups of genes called locus families. We defined all genes in a locus family as orthologues.

**Figure 1:**
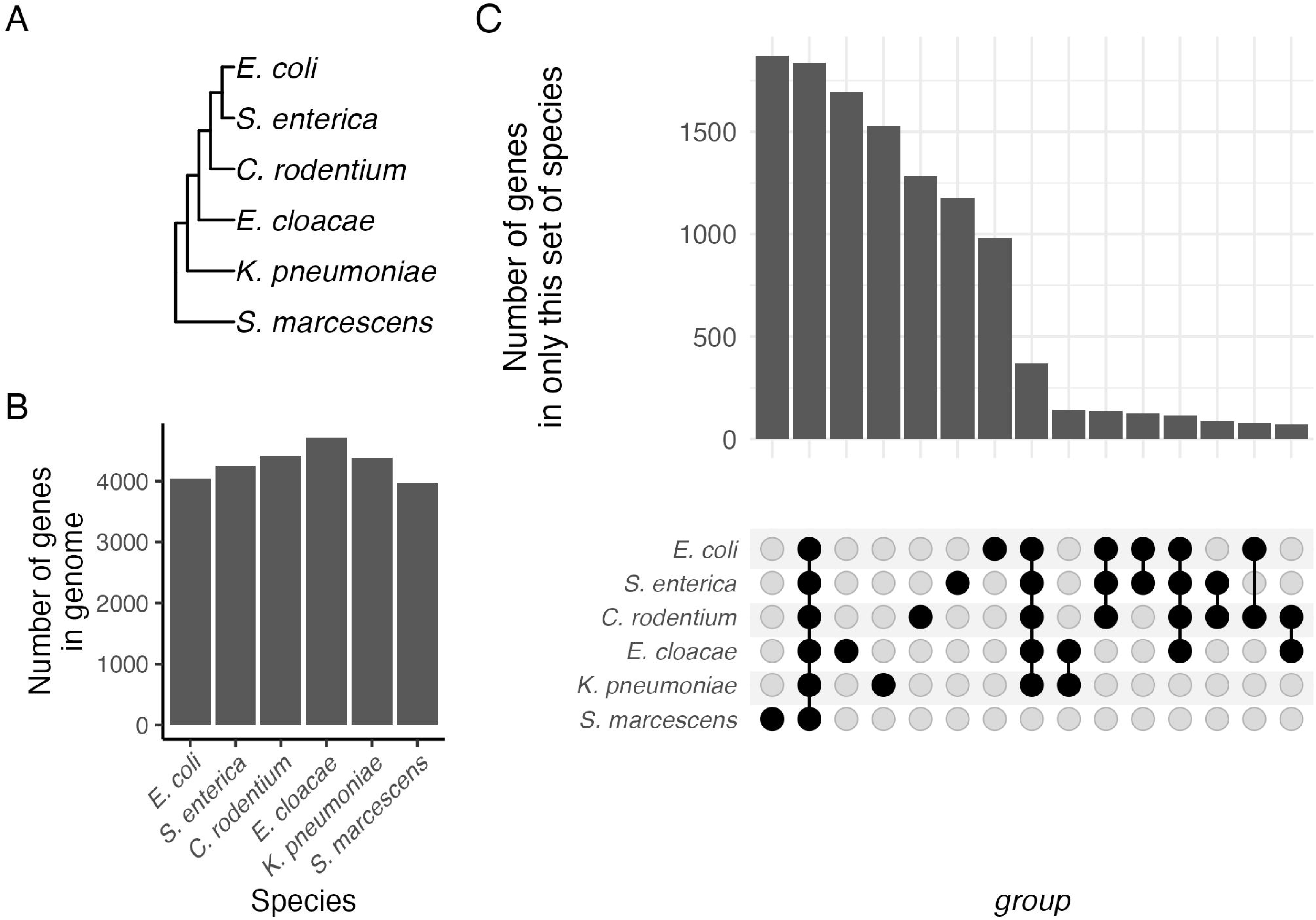
Patterns of shared and unique genome content. (A) Phylogeny of the six species in this study. The phylogeny was constructed by inferring the gene tree for each core gene by maximum likelihood with FastTree (66), followed by reconciliation of the gene trees using ASTRAL (67). The inferred phylogeny is consistent with prior systematic work with this group of organisms (12). (B) The total number of protein coding genes found in the genome of each of the six species. (C) Upset plot of shared and unique genes in these species. Bar heights represent the number of genes found only in this set of species, as indicated by the dots below the bars. For example, the first bar on the left shows the number of genes that are found only in *S. marcescens,* while the second bar represents genes found in all six genomes, the core genome. Only sets with more than 50 genes are shown.

There were 1838 genes found in every genome, a set of genes denoted the core genome. The core genome accounted for 39% to 46% of all genes in a genome. The number of species-specific genes varied from 981 to 1872 (Figure 1C), which accounted for 24% to 47% of all genes in the genome. Most genes in a species were either species-specific or ancestral to the group in this analysis.

### Limited conservation of patterns of differential expression in response to temperature shift

We investigated how the six species responded to a change in temperature from 37°C to 15°C. All species underwent a lag phase after the shift to 15°C, but all were growing again after 3 hours at 15°C (Table S1). After three hours at 15°C, strains were expected to be past the short-term response to cold (involving changes in DNA supercoiling) and into the longer-term response, involving transcriptional changes driven by changing activity of transcription factors (8).

We then performed an RNA-seq experiment on samples growing exponentially at 37°C (immediately before a shift to 15°C), and then after three hours growth at 15°C. Principal components analysis (PCA) confirmed that all samples from each species at each temperature clustered together (Figure 2A), with differences apparent both among species and between the two temperatures. 626 to 1057 genes (14 and 26% of all protein coding genes in the genome) were significantly differentially expressed between the two temperatures across the species (Figure 2B; File S1). (Differential expression was defined as an FDR-adjusted p-value < 0.01 and an absolute log_2_-fold change > 2.) We found little conservation of the set of DE genes in each species (Figure 2C,D). For example, each species had between 274 and 505 genes uniquely DE in that species. In contrast, there were only 16 genes that are DE in every species studied (Table S2). The transcriptional response to low temperature varied strongly among these species.

**Figure 2:**
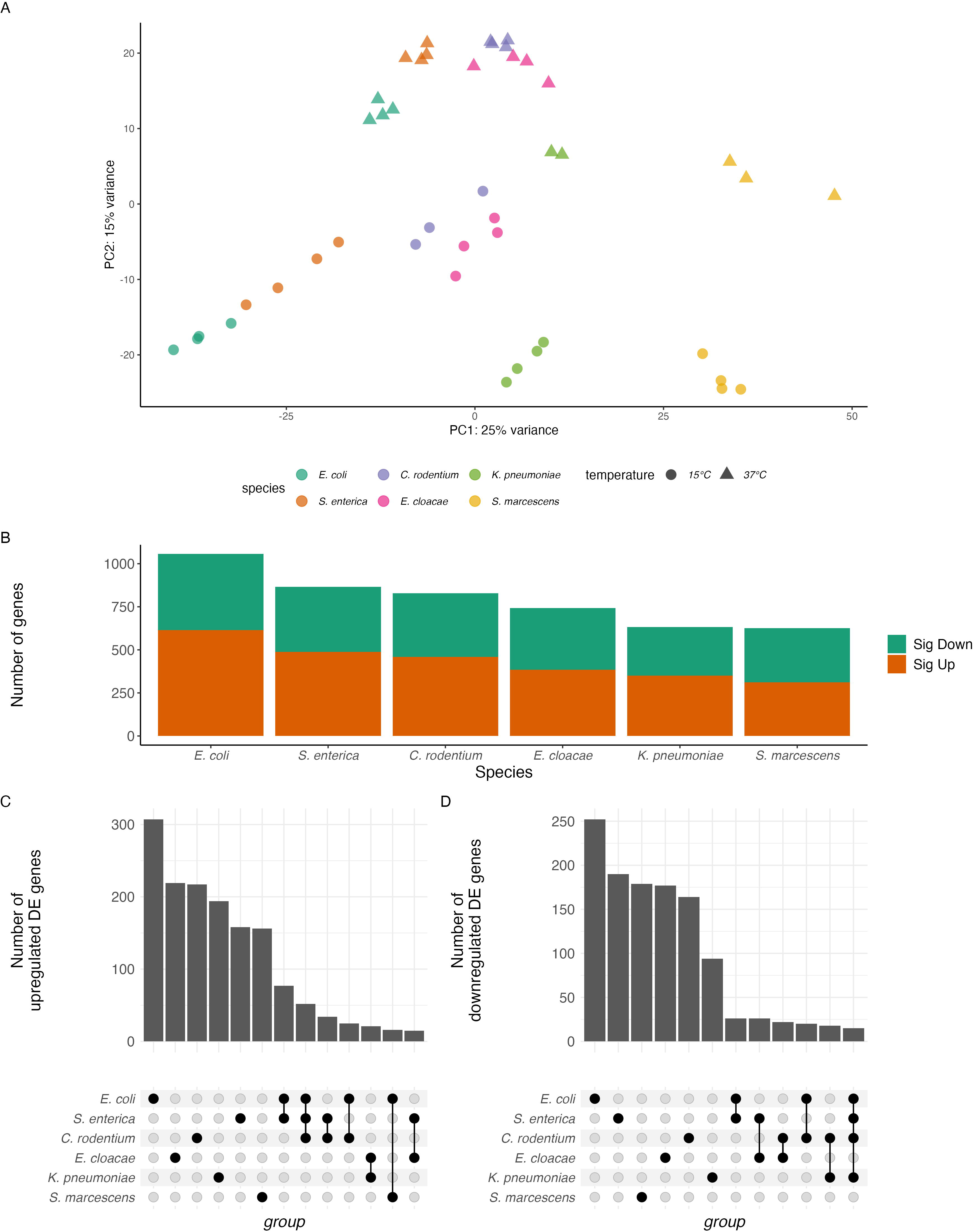
Patterns of differential expression in response to temperature shift of all genes in the genome. (A) Principal component analysis of expression of the 1838 core genes in all six species at 37°C and 15°C. (B) The number of significantly DE genes (FDR-adjusted p < 0.01, absolute log_2_-fold change > 2). Upregulated genes are expressed more highly at 15°C than 37°C. (C, D) Upset plot of shared and unique patterns of DE genes in these species. Bar heights represent the number of genes that are (C) upregulated or (D) downregulated, while the dots below represent the species in that group. Only sets with 15 or more genes are shown.

Each genome contained between 981 to 1872 species-specific genes (Figure 1B), which could be a major contributor to the limited conservation of DE genes. However, these species-specific genes were less likely to be DE between temperatures than expected if all genes had equal probability of being DE (Figure 3A; chi-square test, p < 5 x 10^-4^ for all six species.) In contrast, genes in the core genome were more likely to be DE than expected by chance (Figure 3B; chi-square test, p < 2 x 10^-3^ for all six species). Core genes were particularly likely to be DE in each species, but which core genes were DE in each species was not well-conserved (Figure 3C,D). For example, of the 561 core genes that were DE in *E. coli*, 231 were not DE in any other species, even though all species in this study had those genes. There was not a total lack of conservation of expression pattern, however. For example, there were 60 core genes that were DE in both *E. coli* and *S. enterica* and in no other species. Species-specific expression patterns of core genes strongly contributed to the distinctive transcriptional response to temperature shift of each species.

**Figure 3:**
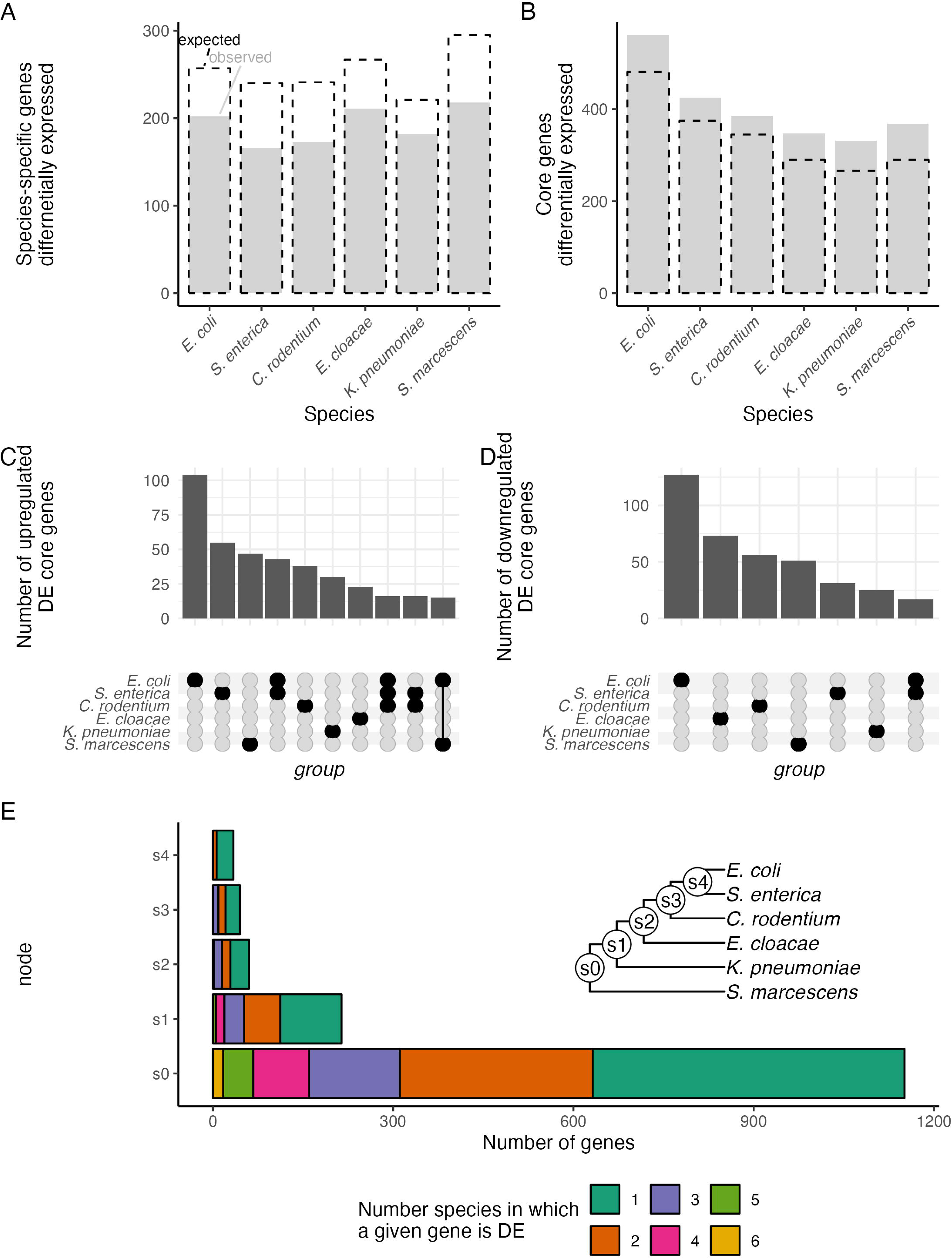
Role of core vs species-specific genes in divergence of transcriptomes. (A, B) The number of DE genes among the (A) species-specific genes and (B) the core genes. Observed counts are shown as gray bars, while the expected counts are shown with black dashed lines. Expected values were calculated as the number of genes in that category multiplied by the proportion of all genes that were DE. In every case, the observed count is significantly different from the expected count (chi-square test, p < 5 x 10^-4^ for each comparison in (A), p < 2 x 10^-3^ for each comparison in (B)). (C, D) The number of significantly DE core genes that are (C) upregulated or (D) downregulated in response to temperature shift. Upregulated genes are expressed more highly at 15°C than 37°C. Bar heights represent the number of genes, while the dots below represent the species in that group. Only sets with 15 or more genes are shown. (E) Distribution of differentially expressed genes among members of a clade. For each gene present in all species descended from a given node, total bar length represents the number of genes that are DE in at least one species. Length of colored segments of the bar represent the number of genes for which that many species are DE. (Inset) Phylogeny of the species, with nodes labels.

We extended our analysis to the other internal nodes on the phylogeny and examined not just the core genes but all those genes that show some level of sharing between species. At each node on the phylogeny, we considered those genes that had an ortholog in each descendent of that node, but in none of the other species, and were DE in at least one of the species. In every case, we found that if a gene was DE in at least one species, the most likely outcome was that it was DE in only that one species (Figure 3E) with decreasing frequencies as the number of species increased. Thus, even when genes were shared between multiple species, patterns of differential expression were primarily not conserved.

Another way to visualize the overall similarity of the transcriptional response or lack thereof is PCA. Principal component 2 of the expression levels of the 1838 core genes, which accounted for 15% of the variance in gene expression, separateed 15°C samples from 37°C samples (Figure 2A). (PC1 and other principal components not visualized in Figure 2A did not clearly separate samples based on temperature.) Thus, there was some conserved difference in the expression of core genes between 15°C and 37°, but it was a small proportion of all variation in gene expression among the species.

### Functions of genes that are DE between warm and cold environments

To better understand the functional consequences of the lack of conservation of DE genes, we used Gene Ontology (GO) enrichment to understand what groups of genes were differentially expressed. We used eggNOG-mapper (24) to assign GO terms to each gene in each genome. (This was done even for genomes like *E. coli* K-12 that had pre-existing GO annotations so that all species had comparable annotations in our analysis.) As a check on the quality of the annotation, we examined the GO terms assigned to each orthologue of the core genome. We found that 1668 of 1838 (91%) of core genes had identical GO annotations in all six genomes, and most of the remaining 118 core genes had identical annotations for 5 of 6 genomes, suggesting that the annotation worked consistently for most core genes.

We then performed GO enrichment analysis using topGO (25) on the RNA-seq data from each species. Consistent with the lack of conservation of differentially expressed genes, there was overall little conservation of enriched GO terms across the genomes (Figure 4; File S2). Of the 151 GO terms that were enriched in at least one species, none were enriched in all six species or in five of the six species, and only 10 were enriched in three or four species (Table S2). 107 of the 151 terms were enriched in only a single species.

**Figure 4.**
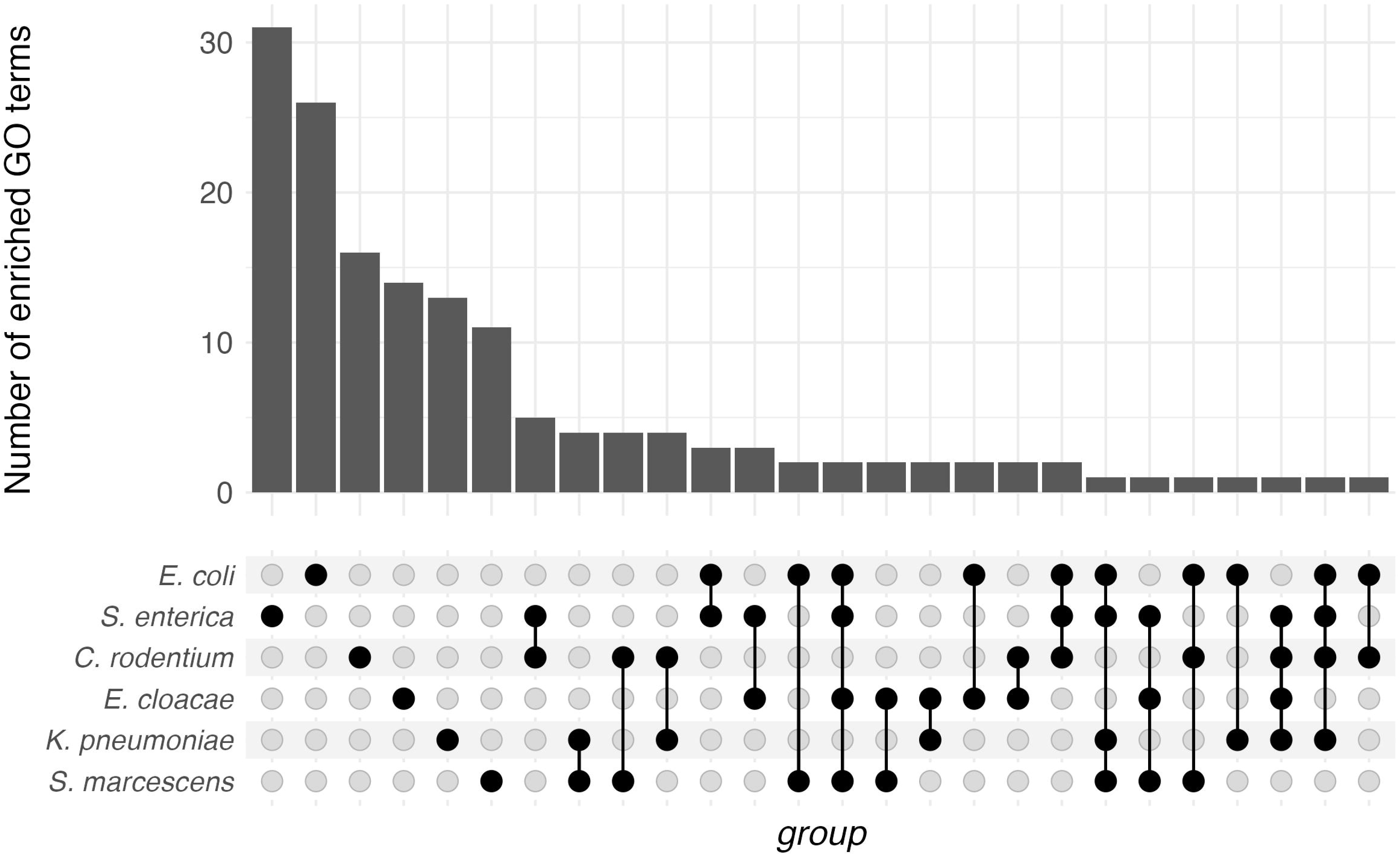
Shared patterns of GO enrichment. Bar heights represent the number of GO terms enriched, while the dots below represent the species involved in that comparison. All observed sets are shown.

### Iron metabolism genes are affected by temperature in multiple species

One group of GO terms that were enriched in multiple genomes were those associated with iron acquisition and use. Enterobactin is a siderophore that is synthesized by many species of Enterobacteraceae (26). This molecule is synthesized inside the cell and secreted into the environment, where it can bind to ferric iron. The iron-enterobactin complex is then actively transported into the cell where the enterobactin is cleaved, freeing the iron (26). Iron is scarce inside of mammalian hosts and genes that allow for iron scavenging are important virulence factors in many of the species in this study (27). The enterobactin biosynthesis term (GO:0009239) was enriched in *E. coli, S. enterica, C. rodentium,*and *K. pneumoniae*, with the associated genes downregulated at 15°C in these species (Figure 5A). Genes for enterobactin transport (GO:0042930) were enriched in *S. enterica, C. rodentium, E. cloacae*and *K. pneumoniae*, with the associated genes also downregulated (Figure 5B). The *E. coli* results are consistent with previous findings that 37°C increases the expression of iron utilization genes in that organism relative to lower temperatures (10). The overall picture is that *S. enterica, C. rodentium,* and *K. pneumoniae*, like *E. coli,* also express genes for iron acquisition less at 15°C than at 37°C. That said, this was not a completely conserved regulatory pattern, as enterobactin biosynthesis levels were largely unchanged in response to temperature in *E. cloacae* and *S. marcescens*.

**Figure 5:**
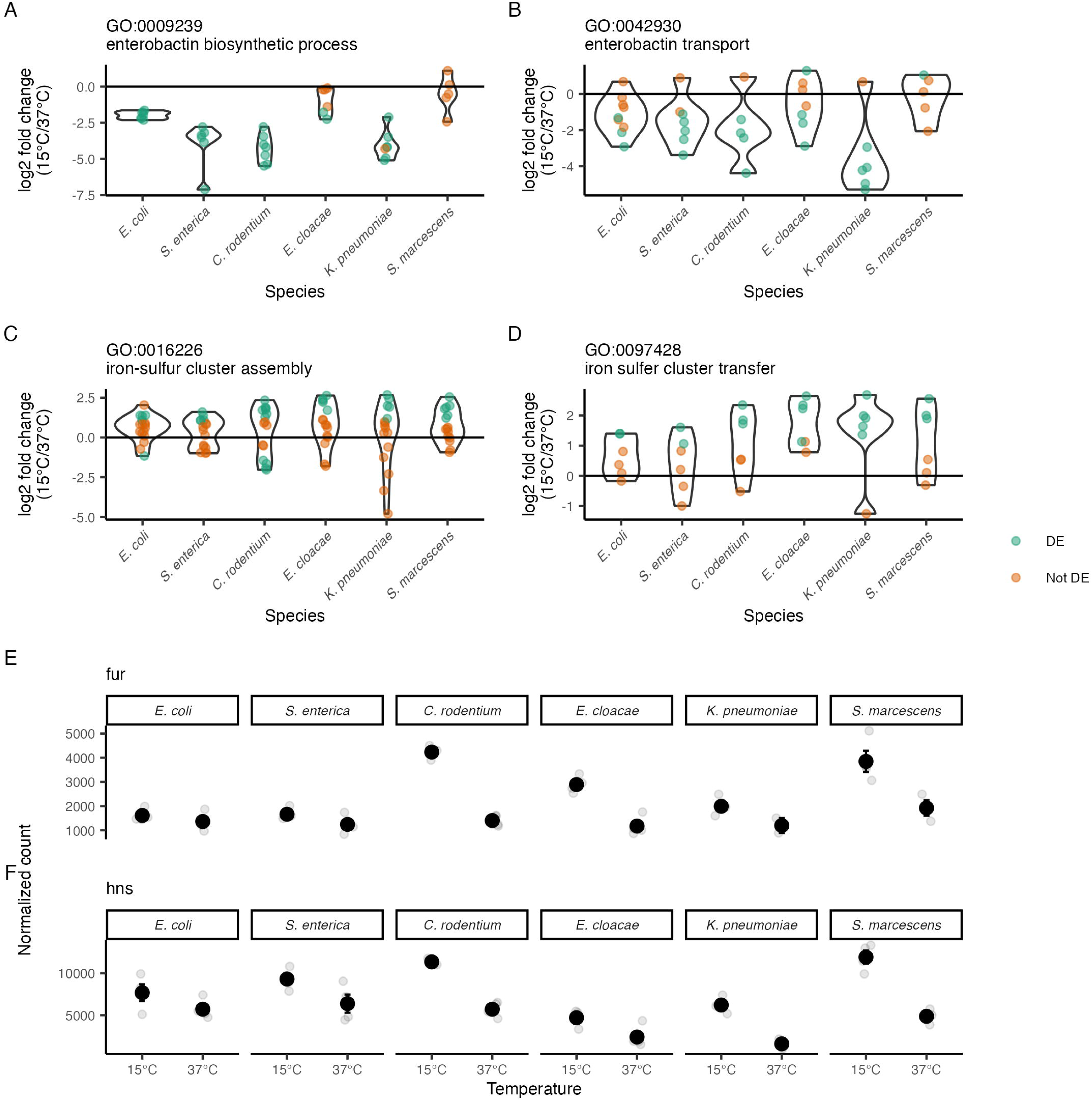
Genes involved in iron metabolism change expression in the cold. (A-D) Log_2_-fold change of gene expression between 15°C and 37°C for genes annotated as involved in several aspects of iron metabolism (A) enterobactin biosynthesis, (B) enterobactin transport, (C) iron-sulfur cluster assembly, and (D) iron-sulfur cluster transfer. Green dots represent a single DE gene, while orange dots are a gene that is not DE. Violin plots show the distributions of log_2_-fold changes of the individual genes. (E, F) The expression of genes coding for two global regulators, *fur* and *hns,* previously identified as associated with these phenotypes. Black dots represent the mean, error bars are the SEM, and gray dots are individual measurements.

One notable use for iron in enteric bacteria is the production of iron-sulfur clusters. Genes associated with iron-sulfur cluster assembly (GO:0016226) that were significantly DE were all upregulated in 15°C relative to 37°C in most species (Figure 5C). However, there were multiple genes expressed at lower levels in *C. rodentium* at 15°C and one such in *E. coli.* Genes associated with the transfer of mature iron-sulfur clusters into proteins (GO:0097428) that were significantly DE were primarily upregulated (Figure 5D), although the number of DE genes varied among the species.

One key regulator of genes associated with iron metabolism in many species is the protein Fur. Fur is primarily a transcriptional repressor (28), but it can serve as a positive regulator in cases where it disrupts repression by another protein, such as its action as an anti-silencer of H-NS at the *ftnA* promoter (29). Fur mRNA levels were significantly higher at 15°C than 37°C only for *C. rodentium* and *E. cloacae* (Figure 5E). *E. coli* and *S. enterica* showed clear repression of enterobactin biosynthesis with non-significant (and very modest) changes in *fur* levels. Temperature regulation of *hns* mRNA also varies across species, with levels significantly higher at 15°C only for *K. pneumoniae* and *S. marcescens* (Figure 5F). Fur protein activity is allosterically regulated by iron (28), so there is not a direct relationship between *fur* mRNA levels and Fur activity. Similarly, H-NS activity is influenced by both physico-chemical factors and by other proteins (30–34), so there is no simple relationship between *hns* mRNA levels and H-NS repression. The effect of increasing *fur* and *hns* mRNA levels at low temperature on the transcription of iron uptake genes remains to be determined.

### Other enriched GO terms

Genes involved in the TCA cycle and with glycine metabolism were significantly enriched in four genomes each (Table S2). (TCA cycle genes are also annotated as involved in citrate metabolism, GO:0006101). These genes were largely upregulated in response to cold (Figure S1A, B), consistent with a previous proteomic study (35). Genes involved in single-species biofilm formation were enriched in three species (Table S2, Figure S1C), replicating previous results from *E. coli* (2), and suggesting that this response is conserved in related species. Genes annotated as being involved in the response to cold were only enriched in *E. coli* and *S. enterica* (Table S2, Figure S1D). Most experimental information used to create GO annotations comes from experiments in *E. coli*, so the lack of enrichment of response to cold genes in more distantly related species is consistent with the lack of conservation in what genes are DE.

### Species-specific gene expression changes in response to temperature

While a few GO terms were enriched across multiple species, 70% of enriched GO terms were species-specific. Here we highlight several species-specific responses. Similar plots of genes associated with a GO term enriched in at least one species are available in the Zenodo archive (DOI <u>10.5281/zenodo.11460468</u>) for interested readers.

### Ribosomes and electron coupled proton transport are enriched only in E. coli

GO terms for ribosome assembly (GO:0000027 and GO:0000028, for ribosomal large and small subunit assembly, respectively) were enriched in the DE genes of *E. coli* but not in other species (Figure S2A). The genes with those GO terms included the structural components of the ribosome as well as factors that modulate ribosome assembly. Those DE ribosomal genes were all downregulated in *E. coli*. Genes coding for proteins that produce the proton motive force via electron transport (GO:0015990) were also enriched in *E. coli* but not the other species (Figure S2B). Those genes, which were all downregulated, were *cyoABCD*, and *nuoLMN,* encoding the cytochrome bo3 quinol:oxygen oxidoreductase and NADH:ubiquinone oxidoreductase I respectively. The mechanism and consequences of this *E. coli-*specific regulation remains to be determined.

### Colanic acid biosynthesis is enriched only in C. rodentium

Colanic acid is the capsular polysaccharide in *E. coli* and some relatives (36–38). Genes annotated with the GO term for colanic acid biosynthesis (“GO:0009242”) were enriched in *C. rodentium* but not in other species. Some genes in the *wca* cluster, which encodes the biosynthetic enzymes for colanic acid (37), were not annotated with this GO term, so we created a custom gene set that included all genes annotated with GO:0009242 plus all genes from the *wca* cluster that were missing that annotation. (The strains of *K. pneumoniae* and *S. marcescens* we worked with do not have orthologues to the *wca* genes found in the other four species.) 70% of genes in the combined set were significantly upregulated in *C. rodentium*, while many fewer (if any) genes were upregulated in each of the other species (Fig 6A).

**Figure 6:**
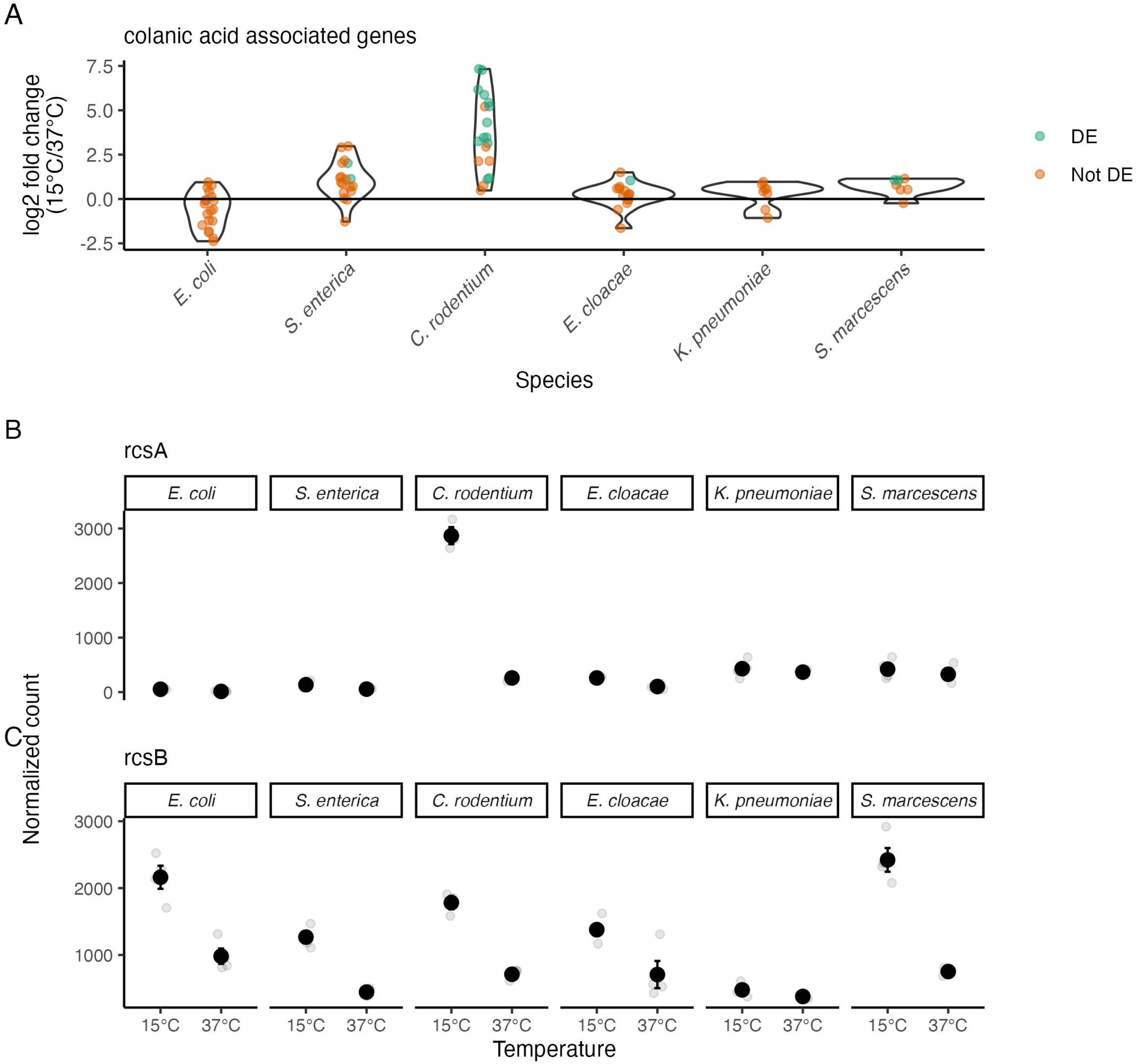
Upregulation of colanic acid related genes only in. C. rodentium. (A) Log_2_-fold change of gene expression between 15°C and 37°C for genes associated with production and use of colanic acid. Green dots represent a single DE gene, while orange dots are a gene that is not DE. Violin plots show the distributions of log_2_-fold changes of the individual genes. (B, C) Expression of two known regulators of this process, *rcsA* and *rcsB*. Black dots represent the mean, error bars are the SEM, and gray dots are individual measurements.

In *E. coli*, the Rcs phosphorelay system is an important regulator of colanic acid biosynthesis (39). The response regulator RcsB forms a heterodimer with the RcsA protein, and these two proteins are required for maximal capsule expression in that organism (40). The gene *rcsA* had an expression pattern that closely matched the pattern of its target genes, being significantly upregulated 10-fold in *C. rodentium,* while being upregulated at most 2-fold in the other five species (Fig 6B). In contrast, RcsB showed a similar pattern of expression across five of the six species, rather than being upregulated only in *C. rodentium* (Fig 6C). The evolution of species-specific regulation of RcsA in *C. rodentium* is a promising candidate for the evolutionary change that brought about the species-specific pattern of the colanic acid biosynthesis genes.

### RpoS regulation of the response to temperature shift

The alternative sigma factor RpoS is known to be activated at low temperature in *E. coli* where it regulates the expression of several hundred genes (41). We compared the role of RpoS in *E. coli* and its close relative *S. enterica* during growth at 15°C. Western blotting demonstrated that RpoS had a similar pattern of expression in both species. RpoS could accumulate during exponential growth at 15°C to a similar level as in stationary phase at 37°C, but not during exponential growth at 37°C (Figure S3).

RNA-seq experiments with Δ*rpoS* strains of *E. coli* and *S. enterica* allowed us to define the RpoS regulon at 15°C. During exponential growth at 15°C, 232 genes were DE between wild-type and Δ*rpoS E. coli.* This represented 22% of the number of genes that were DE between 37°C and 15°C in the wild-type strain. Reassuringly, of the 232 DE genes in *E. coli* in this study, 189 (81%) were previously identified as RpoS-regulated at 15°C by Adams *et al.* (41). In *S. enterica,* 189 genes were DE between the wild-type and Δ*rpoS* strains, 80% of the number of genes that are RpoS-regulated in *E. coli.* RpoS regulates many genes in the cold, although only a fraction of all the genes that are cold-regulated.

The role of RpoS in regulating the 15°C response is similar but distinct in *E. coli* and *S. enterica.* Principal component analysis of the expression of core genes clearly separated the two species, but also found the same direction of change in PCA space between wild-type and Δ*rpoS* strains of both species (Figure 7A). 83 genes were DE in both *E. coli* and *S. enterica* (Figure 7B). This included genes previously reported to be RpoS-dependent (42–44) such as *dps, katE, mlrA, osmC, osmY, otsA, otsB, sodC,* and *wrbA.* Consequently, GO terms involved in response to osmotic stress or response to hydrogen peroxide were enriched in both species. Most of the RpoS-regulon in *E. coli* or *S. enterica* was not composed of the 83 genes that were RpoS-regulated in the other species (Figure 7B). For example, of the 232 RpoS-regulated genes in *E. coli,* 149 of those genes (64%) were not RpoS-regulated in *S. enterica.* Those 149 genes were composed of 50 genes that were not present in *S. enterica,* and 99 genes that had an ortholog in *S. enterica*, but that ortholog was not RpoS-regulated in *S. enterica.* Like the entire cold-response regulon, divergence of the RpoS-regulon between *E. coli* and *S. enterica* was due to a mixture of both different regulation of genes shared between the two species and the presence of species-specific genes.

**Figure 7:**
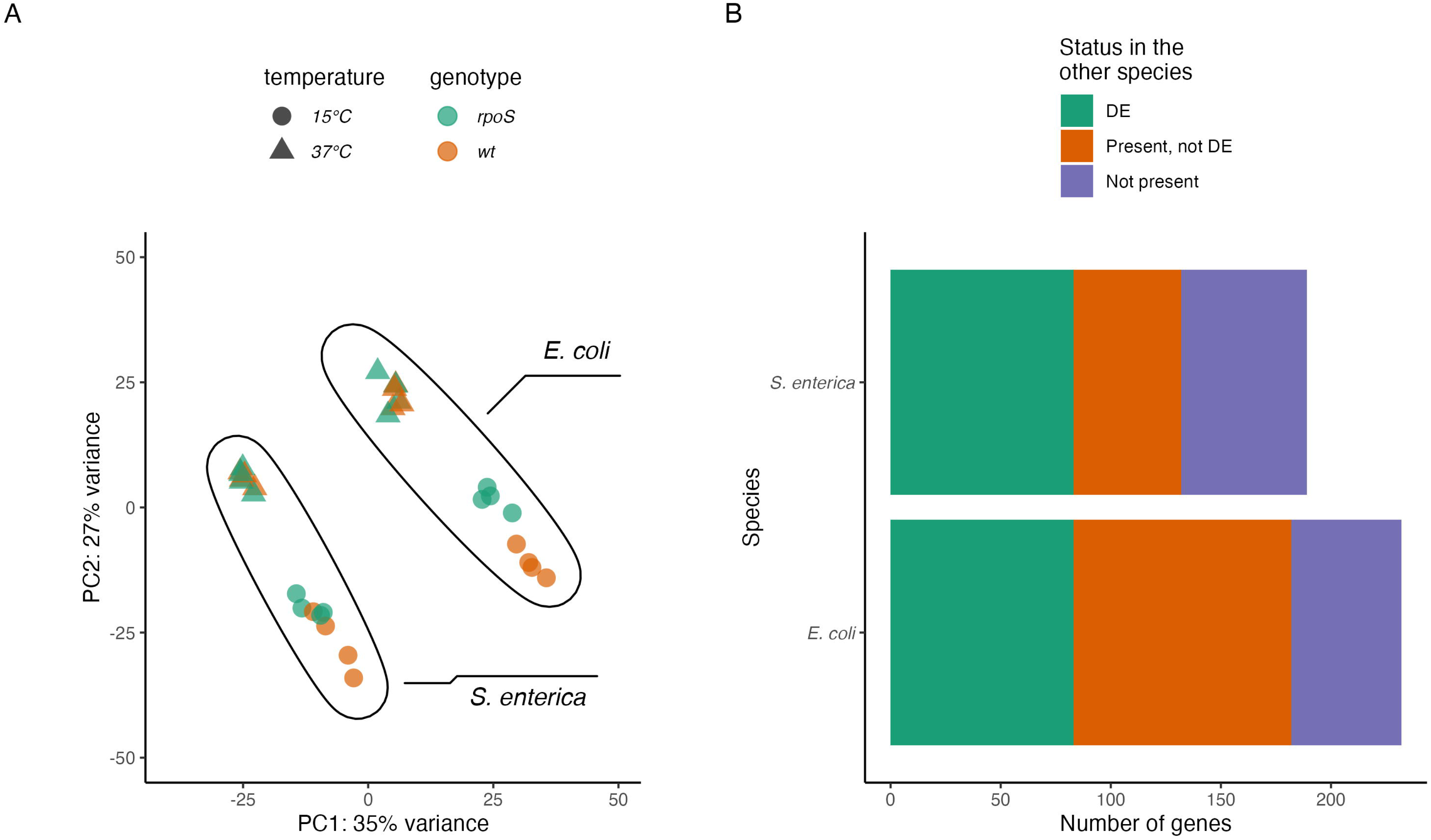
Partial overlap of the RpoS regulon of. E. coli **and** S. enterica **at 15°C.** (A) Principal component analysis of expression of the 1838 core genes in wild-type and Δ*rpoS E. coli* and *S. enterica* at 37°C and 15°C. (B) Size of the RpoS regulon in *E. coli* and *S. enterica.* Bar lengths are the number of RpoS-regulated genes in that species, with colors denoting the status of those genes in the other species. For example, the orange part of the *E. coli* bar represents the 99 genes that are RpoS-regulated in *E. coli* that are also present in *S. enterica* but are not RpoS-regulated in *S. enterica*.

Genes that were RpoS-regulated only in *E. coli* include *astABCDE,* responsible for arginine catabolism (GO:0006527), *dppABCDF*, involved in di-peptide transport, *gabD*, *astC*, and *ldcC*, involved in lysine metabolism (GO:0006553). These genes have orthologs in *S. enterica* but were not significantly DE in that organism, pointing to a role for RpoS in amino-acid catabolism in *E. coli* that is not present in *S. enterica.* RpoS also had an *E. coli-*specific role in regulation of genes *gadE, gadW, gadX, hdeA, hdeB,* and *slp,* which are part of the acid fitness island not present in related species like *S. enterica* (45).

Genes that were RpoS-regulated only in *S. enterica* include *glgB, glgC, glgP,* and *glgX*, responsible for the biosynthesis and catabolism of glycogen. (These genes are expressed more highly at 15°C than 37°C in both *E. coli* and *S. enterica* but are only RpoS-regulated in *S. enterica*.) There were also genes that were RpoS-dependent and present only in *S. enterica*, like *invA*, *invE*, *prgI, sipD*, and *sprB*. These genes are all located on the Salmonella Pathogenicity Island 1 and are not present in *E. coli* (46). Much like the broader response to low temperature, the RpoS-regulon at 15°C was mostly not conserved between *E. coli* and *Salmonella* due a mix of species-specific genes and different regulation of conserved genes.

## Discussion

All the species studied in these experiments responded to a shift in growth temperature from 37°C to 15°C by changing the expression of a large proportion of their genes; the number ranged from 626 to 1057 genes, or 14 to 26% of all genes in the genome. For *E. coli*, this was more than twice the number of genes than were previously reported in microarray studies of a shift from 37°C to 23°C (2, 10). The difference can probably be attributed to the combination of the more extreme temperature shift in this study, and the difference in technologies used to measure mRNA abundance.

A major finding of this study is the lack of conservation of the transcriptional response to low temperature across the *Enterobacteriaceae*. All the species grew markedly slower at 15°C than at 37°C and all changed expression of hundreds of genes in response to the temperature shift. However, the identity of these genes was mostly not conserved among species. The lack of conservation of which genes were DE is due primarily to species-specific transcriptional patterns of shared genes. While species-specific genes did contribute to the diversity of differentially expressed genes in this group, these species-specific genes were less likely to be DE than was the core genome. Horizontal gene transfer is the major way that bacterial genomes become different from other each other, but the transcriptomes in this study mostly became different by species-specific patterns of expression of shared genes.

*E. coli* responded to low temperature by changing the expression of more of its genes than any of the other species studied here. It is possible that the K-12 strain of *E. coli* is more lab-adapted than the other strains used in this study (47, 48) and so responds more strongly to low temperature than the other strains. (The history of some of the strains used in this study is not well documented.) It is also possible that K-12 strains are typical of all *E. coli* in the number of genes differentially regulated by temperature shift, or that some of the other strains in this study are just as lab-adapted as *E. coli* K-12. Now that we have documented extensive transcriptional divergence using single strains of each species, future experiments should examine multiple strains sampled from the diversity of a single species to determine the extent of intraspecific variation in transcriptomic response to low temperature.

The lack of conservation of which genes were differentially expressed extends to a lack of conservation of phenotypes as predicted by enriched GO terms. Most of the GO terms enriched in a species were not shared with other species, which implies that each species in this study may have different phenotypic responses to the same stress. For example, only *E. coli* responded to the lower temperature by down-regulating the mRNAs of ribosomal proteins. It is well known that the abundance of ribosomal proteins in *E. coli* is lower at slower growth rates (49), and our data on the abundance of ribosomal protein mRNAs are consistent with this finding. The other species in this study all grew more slowly at 15°C than at 37°C, and some grew as slowly as *E. coli* at 15°C, but none showed the same down-regulation of ribosomal protein mRNA. It is possible that these other species do not achieve down-regulation of ribosomal activity at lower growth rate via the same mechanism, or do not down-regulate ribosomal activity at all.

The alternative sigma factor RpoS is one regulatory protein that plays a role in the transcriptional response to temperature shift. Much like the overall cold response, the RpoS-dependent part of the cold response was only partially conserved between *E. coli* and *S. enterica.* The divergence between the two species was due to a mix of divergent expression of genes shared between the two species and species-specific genes that are part of the RpoS-regulon. The divergence in RpoS-dependence of genes shared by *E. coli* and *S. enterica* was somewhat surprising given that both species produce similar amounts of RpoS at 15°C. Divergence in RpoS-dependence of shared genes could be due to changes in the RpoS protein itself, in the promoters of these divergently expressed genes, or could be due to changes in the many trans-acting factors that interact with RpoS to determine the transcriptional output of a promoter (44). Understanding this molecular basis of these changes is an important future area of research.

In other systems, species-specific genes are a substantial part of the divergence of regulons between species pairs. For example, the OmpR protein regulates transcription of a group of genes in response to physical stresses like osmolarity and pH. The OmpR regulon in *E. coli* and *S. enterica* is only weakly conserved, and the authors noted that many species-specific genes were in the regulon (50). The RovA protein regulates the transcription of important virulence genes in *Yersinia*. The RovA-regulon of *Yersinia enterocolitica*and *Y. pestis* is also only weakly conserved due at least in part to the species-specific genes regulated by RovA in each species (51). The transcriptional regulator YhaJ regulates different sets of genes in strains of enterohemorrhagic and uropathogenic *E. coli*, some of which are strain-specific (52, 53). This paper extends the observation of regulon divergence to the RpoS-regulon and shows that while species-specific genes do have a role in divergence of the regulon, shared genes are a disproportionate part of the species-specific regulons.

In conclusion, we studied the transcriptional response to a change in temperature from 37°C to 15°C in five species of the *Enterobacteriaceae*and one outgroup. We found that there was limited conservation of this response, with each species changing the transcription of hundreds of genes that are unchanged in the other species. This divergence was not due primarily to species-specific genes in each genome, as core genes were more likely than expected to be differentially expressed in response to temperature change. This pattern of evolutionary divergence was also true of RpoS-regulon, where divergence was due more to shared genes being RpoS-regulated in only a single species than it was due to species-specific genes being part of the regulon. These results highlight the importance of the evolution of transcriptional patterns of shared genes in the evolution of bacterial responses to stress.

## Materials and Methods

### Strains and strain construction

Strains of six species were used in this study: *E. coli* K-12*, S. enterica* sv. Typhimurium 14028s*, C. rodentium* DBS100*, E. cloacae* ATCC13047*, Klebsiella pneumoniae*MKP103, and *Serratia marcescens* Db11. Strains with wild-type alleles for *rpoS* were used for each species. In addition, strains of E. coli and S. enterica with existing Δ*rpoS* alleles were used (Table S3). To construct the isogenic Δ*rpoS* strain of *S. enterica*, the Δ*rpoS::cm* allele of a *S. enterica* 14028s strain provided by the McClelland Lab (54) was moved by P22 transduction (55) into our lab isolate of *S. enterica* 14028s.

### Growth conditions

Strains were inoculated from lll80°C frozen cultures and grown aerobically overnight in 5 mL of Luria-Bertani (LB) broth (1% tryptone, 0.5% yeast extract, 1% NaCl) at 37°C with shaking at 225 rpm. The next morning, 250μL of each culture was diluted into 25 mL of LB in a 250 mL borosilicate flask and grown at 37°C in an incubator shaking at 225 rpm. When cultures reached OD_600_ of 0.25 - 0.35, flasks were transferred to a water bath at 15°C shaking at 225 rpm. (OD_600_ of 0.3 is in the middle of exponential growth in these cultures.) When cultures were grown for RNA and protein isolation, no measurements of OD_600_ were performed at intermediate time points between the shift and sampling of RNA and protein. To minimize confounding factors, we always grew all 8 strains in parallel for each experimental replicate.

### Measurement of growth parameters

When cultures were grown to measure growth rate, the growth conditions mentioned above were used, except that OD_600_ measurements were taken approximately every 15 minutes in the 37°C portion of the experiment (from the start of the experiment until an OD_600_ of 0.25 – 0.35 was reached), and every 20 to 30 minutes after the shift to 15°C. (Cultures reached maximum OD_600_ values of less than 1 after three hours at 15°C.) Growth rates at 37°C and 15°C, as well as the length of the lag phase after the shift to 15°C were estimated using the “Growth Rates Made Easy” method of Hall *et al.* (56), as implemented in the R package growthrates (57). To minimize confounding factors, all 8 strains were always grown in parallel for each experimental replicate.

### RNA Isolation

For RNA isolation, 1.5 mL of bacterial culture was pelleted and resuspended in 500 μL of Trizol pre-heated to 65°C and then stored at −80°C until use. Samples were purified on a column using the Zymo Direct-Zol RNA Miniprep kit. They were then treated twice with DNase I using Turbo DNA-free (Invitrogen) for 30 minutes each at 37°C to ensure that all DNA was degraded. The samples were further purified on a column using the Zymo RNA Clean and Concentrator Kit and again stored at −80°C until sequencing. The concentration of the samples was checked using a NanoDrop spectrophotometer and degradation was checked for by electrophoresing samples on an ethidium bromide agarose gel. Four biological replicate samples were collected for each strain at each temperature.

### Generation of RNA-Seq data

Illumina cDNA libraries were generated using a modified version of the RNAtag-seq protocol (58), which creates tagged pools of RNA samples for cDNA synthesis and sequencing. Briefly, 500 ng to 1 μg of total RNA was fragmented by heating to 94°C in NEB Alkaline Phosphatase Buffer, depleted of any residual genomic DNA, dephosphorylated, and ligated to DNA adapters carrying 5’-AN_8_-3’ barcodes of known sequence with a 5’ phosphate and a 3’ blocking group. Barcoded RNAs were pooled and depleted of rRNA using the RiboZero rRNA depletion kit (Epicentre). Pools of barcoded RNAs were converted to Illumina cDNA libraries in 2 main steps: (i) reverse transcription of the RNA using a primer designed to the constant region of the barcoded adaptor with addition of an adapter to the 3’ end of the cDNA by template switching using SMARTScribe (Clontech) as described (59); (ii) PCR amplification using primers whose 5’ ends target the constant regions of the 3’ or 5’ adaptors and whose 3’ ends contain the full Illumina P5 or P7 sequences. cDNA libraries were sequenced on the Illumina [NextSeq 500] platform at the Broad Institute to generate paired end reads. Each RNA pool consisted of 16 samples, a 15°C and a 37°C sample from each of the 8 strains. Four total pools were constructed and sequenced.

### RNA-seq Data Analysis

Sequencing reads from each sample in a pool were demultiplexed based on their associated barcode sequence using custom scripts. Up to 1 mismatch in the barcode was allowed provided it did not make assignment of the read to a different barcode possible. Barcode sequences were removed from the first read as were terminal G’s from the second read that may have been added by SMARTScribe during template switching.

Reads were aligned to the reference genomes (Table S1) using BWA v0.7.17 (60). Reads were counted with htseq-count v2.0.1 (61). Further analysis was done in R v4.4.0 (62). Plots were created with ggplot2 3.5.1 (63).

Before conducting the analysis reported in this paper, initial quality control was performed to ensure that each sample had at least 1,000,000 read-pairs that mapped to protein coding sequences. One sample below this threshold was removed. In addition, three samples did not cluster with any of the other samples in PCA or a heatmap and were also removed. The samples removed were one from *C. rodentium* at 15°C, two from *K. pneumoniae* at 37°C, and one from *S. marcescens* at 37°C.

Differential expression analysis was performed using the package DESeq2 v1.40.2 (64), with the analysis of each species occurring separately. Genes were considered significant if they had an FDR-adjusted p-value of less than 0.01 and an absolute value of fold-change greater than 2. Readers interested in repeating these analyses with different p-values and fold-change can use the data and scripts in the Zenodo archive. Upset plots were created with the package ComplexUpset v 1.3.3 in R (65). For the comparison of expression levels across species in plots of individual genes, all samples from all species were normalized together as one data set.

eggNOG-mapper (24) was used to assign GO terms to each gene in each genome. GO enrichment for each species was performed with the R package topGO (25). The GO tree for each species was pruned to remove terms with fewer than 5 genes. The weight01 algorithm was used with Fisher’s exact test to calculate p-values on enrichment of GO terms. GO terms were considered significantly enriched if they had a p-value of less than 0.05.

### Protein Isolation & Western Blotting

Total protein was isolated from the same cultures at the same time as RNA samples. Samples with cell density equivalent to 1mL of OD_600_ = 0.3 were pelleted by centrifugation at least 20,000 x g for 30 s, resuspended in 100μL of 2x Laemmli sample buffer (LiCor), heated at 95°C for 5 minutes, and then stored at −20°C until they were used.

A quantitative western blot was used to measure the levels of RpoS in the cell at each time point, in a procedure slightly modified from (15). 10μL of each protein sample was electrophoresed on a 4 - 20% gradient polyacrylamide gel (Bio-Rad) in Tris-glycine running buffer (25 mM Tris base, 250 mM glycine, 0.5% SDS) at 200V for 35 minutes at room temperature. Electrophoresis was used to transfer proteins to a Immobilon-FL PVDF membrane in transfer buffer (48 mM Tris base, 39 mM glycine, 0.0375% SDS, 20% methanol) at 100V for 45 minutes at 4°C. To quantify the total amount of protein in the samples, the membranes were stained with 5 mL of REVERT Total Protein Stain solution and imaged using the LiCor Odyssey CLx imager. The membranes were blocked with 10mL of Odyssey Blocking Buffer for 1 hour at room temperature.

RpoS was detected using an anti-RpoS monoclonal antibody (clone 1RS1, BioLegend). The blocked membranes were labeled with primary antibody (0.4 μg/mL mouse anti-RpoS, 10mL Odyssey Blocking Buffer, 0.2% Tween 20) at 4°C with shaking at 55 rpm overnight. Once labeled, the membrane was washed four times with 10 - 15mL of 1x Tris-buffered saline with Tween 20 for 5 minutes each wash. The membranes were probed with a fluorescent secondary antibody (IRDye 800CW goat anti-mouse, LiCor) that was diluted in 10mL of Odyssey Blocking Buffer with 0.2% Tween 20 and 0.01% SDS. The membranes were incubated with the diluted secondary antibody for 1 hour at room temperature and were washed as described previously, with an additional wash at the end with 1x Tris-buffered saline without Tween 20. Membranes were dried for 2-3 hours between 2 sheets of Whatmann 3MM blotting paper before imaging on a Li-Cor Odyssey CLx imager. RpoS levels were quantified by analyzing the amount of fluorescence of each band with Image Studio v 2.1 software.

Data analysis was performed in R v4.4.0 (62). RpoS levels were normalized by dividing them by the total amount of protein at each time point, as quantified by the total protein stain.

### xenoGI analysis

The program xenoGI v3.1.1 (23) was used to reconstruct the evolutionary history of the genomes used in this study. The phylogeny was inferred using the built-in helper function of xenoGI. This function built a gene tree for each core gene using FastTree 2.1.11 (66), then reconciled all of the gene trees into a species tree using Astral v5.6.3 (67).

## Data Availability

The RNA-seq FASTQ files are deposited in the Gene Expression Omnibus with ascension number GSE267531.

All other data and scripts used for data analysis are in a Zenodo archive at DOI 10.5281/zenodo.11460468

## Acknowledgements

RNA-Seq libraries were constructed and sequenced at the Broad Institute of MIT and Harvard by the Microbial ‘Omics Core and Genomics Platform, respectively. The Microbial ‘Omics Core also provided guidance on experimental design and performed the demultiplexing of RNA-Seq data. Thanks to Michael McClelland, Colin Manoil, the Salmonella Genetic Stock Center, and the *C. elegans* Stock Center for sharing strains, to Eliot Bush for help with xenoGI, and to Danae Schulz and Chris White-Ziegler and anonymous peer-reviewers for feedback and insights.

This material is based upon work supported by the National Science Foundation under Grant No. 1716794.

