## Supplemental Tables S1-S3 and Supplemental Figures S1-S3 for "The transcriptional response to low temperature is weakly conserved across the *Enterobacteriaceae*"

### Supplementary information for Hoang and Stoebel

**Table S1:** Growth parameters at 15°C and 37°C in LB[†]

| Strain | Doubling time at 37°C (mins) | Lag time after shift to 15 °C (mins) | Doubling time at 15°C (mins) |
|---|---|---|---|
| *E. coli* wt | 28.5 ± 0.3 | 63.7 ± 12.2 | 328.0 ± 15.2 |
| *E. coli ΔrpoS* | 25.5 ± 0.2 | 98.1 ± 3.6 | 303.5 ± 6.4 |
| *S. enterica* wt | 29.6 ± 1.7 | 96.2 ± 6.7 | 175.5 ± 3.8 |
| *S. enterica ΔrpoS* | 27.5 ± 1.2 | 97.8 ± 10.1 | 188.8 ± 7.4 |
| *C. rodentium* wt | 28.1 ± 0.6 | 104.0 ± 23.1 | 342.2 ± 30.9 |
| *E. cloacae* wt | 28.0 ± 3.6 | 90.3 ± 29.7 | 394.2 ± 44.6 |
| *K. pneumoniae* wt | 26.8 ± 0.7 | 119.4 ± 7.7 | 268.9 ± 19.8 |
| *S. marcescens* wt | 29.9 ± 1.3 | 123.8 ± 9.8 | 160.9 ± 7.1 |

[†] Values are the mean ± standard error of the mean. N = 3 for all measurements.

**Table S2: Genes differentially expressed in all six species**

| *E. coli* gene name | Direction of regulation |
|---|---|
| *yqjD* | Upregulated (higher at 15°C than 37°C) |
| *ygaM* | Upregulated (higher at 15°C than 37°C) |
| *iscA* | Upregulated (higher at 15°C than 37°C) |
| *gltA* | Upregulated (higher at 15°C than 37°C) |
| *iscR* | Upregulated (higher at 15°C than 37°C) |
| *gcvT* | Upregulated (higher at 15°C than 37°C) |
| *zapC* | Upregulated (higher at 15°C than 37°C) |
| *bssS* | Upregulated (higher at 15°C than 37°C) |
| *rof* | Upregulated (higher at 15°C than 37°C) |
| *pstS* | Upregulated (higher at 15°C than 37°C) |
| *brnQ* | Downregulated (lower at 15°C than 37°C) |
| *mnmC* | Downregulated (lower at 15°C than 37°C) |
| *recO* | Downregulated (lower at 15°C than 37°C) |
| *fhuB* | Downregulated (lower at 15°C than 37°C) |
| *feoB* | Downregulated (lower at 15°C than 37°C) |
| *yidD* | Downregulated (lower at 15°C than 37°C) |

**Table S3:** GO terms enriched in at least one genome

| GO ID | term | number of genomes enriched | *E. coli* | *S. enterica* | *C. rodentium* | *E. cloacae* | *K. pneumoniae* | *S. marcescens* |
|---|---|---|---|---|---|---|---|---|
| GO:0009239 | enterobactin biosynthetic process | 4 | Enriched | Enriched | Enriched | Not Enriched | Enriched | Not Enriched |
| GO:0042930 | enterobactin transport | 4 | Not Enriched | Enriched | Enriched | Enriched | Enriched | Not Enriched |
| GO:0006101 | citrate metabolic process | 4 | Enriched | Enriched | Not Enriched | Enriched | Not Enriched | Enriched |
| GO:0006099 | tricarboxylic acid cycle | 4 | Enriched | Enriched | Not Enriched | Enriched | Not Enriched | Enriched |
| GO:0006546 | glycine catabolic process | 4 | Enriched | Enriched | Not Enriched | Not Enriched | Enriched | Enriched |
| GO:0051641 | cellular localization | 3 | Enriched | Enriched | Enriched | Not Enriched | Not Enriched | Not Enriched |
| GO:0044011 | single-species biofilm formation on inan... | 3 | Enriched | Not Enriched | Enriched | Not Enriched | Not Enriched | Enriched |
| GO:0006979 | response to oxidative stress | 3 | Enriched | Enriched | Enriched | Not Enriched | Not Enriched | Not Enriched |
| GO:0016999 | antibiotic metabolic process | 3 | Not Enriched | Enriched | Not Enriched | Enriched | Not Enriched | Enriched |
| GO:0009312 | oligosaccharide biosynthetic process | 3 | Not Enriched | Enriched | Not Enriched | Not Enriched | Enriched | Enriched |
| GO:0051289 | protein homotetramerization | 2 | Not Enriched | Not Enriched | Enriched | Enriched | Not Enriched | Not Enriched |
| GO:0033212 | iron import into cell | 2 | Not Enriched | Enriched | Enriched | Not Enriched | Not Enriched | Not Enriched |
| GO:0005991 | trehalose metabolic process | 2 | Not Enriched | Enriched | Enriched | Not Enriched | Not Enriched | Not Enriched |

| | | | | | | | | |
|---|---|---|---|---|---|---|---|---|
| GO:0040011 | locomotion | 2 | Not Enriched | Not Enriched | Enriched | Enriched | Not Enriched | Not Enriched |
| GO:0009252 | peptidoglycan biosynthetic process | 2 | Not Enriched | Not Enriched | Enriched | Not Enriched | Enriched | Not Enriched |
| GO:0006974 | cellular response to DNA damage stimulus | 2 | Not Enriched | Enriched | Enriched | Not Enriched | Not Enriched | Not Enriched |
| GO:0006972 | hyperosmotic response | 2 | Not Enriched | Enriched | Enriched | Not Enriched | Not Enriched | Not Enriched |
| GO:0034755 | iron ion transmembrane transport | 2 | Not Enriched | Not Enriched | Enriched | Not Enriched | Enriched | Not Enriched |
| GO:0008360 | regulation of cell shape | 2 | Not Enriched | Not Enriched | Enriched | Not Enriched | Enriched | Not Enriched |
| GO:0006970 | response to osmotic stress | 2 | Not Enriched | Not Enriched | Enriched | Not Enriched | Not Enriched | Enriched |
| GO:0006793 | phosphorus metabolic process | 2 | Not Enriched | Not Enriched | Enriched | Not Enriched | Not Enriched | Enriched |
| GO:0009225 | nucleotide-sugar metabolic process | 2 | Not Enriched | Not Enriched | Enriched | Not Enriched | Not Enriched | Enriched |
| GO:0015718 | monocarboxylic acid transport | 2 | Not Enriched | Enriched | Enriched | Not Enriched | Not Enriched | Not Enriched |
| GO:0009314 | response to radiation | 2 | Not Enriched | Not Enriched | Enriched | Not Enriched | Not Enriched | Enriched |
| GO:0009061 | anaerobic respiration | 2 | Not Enriched | Not Enriched | Not Enriched | Enriched | Not Enriched | Enriched |
| GO:0022900 | electron transport chain | 2 | Not Enriched | Not Enriched | Not Enriched | Enriched | Not Enriched | Enriched |
| GO:0015685 | ferric-enterobactin import into cell | 2 | Not Enriched | Enriched | Not Enriched | Enriched | Not Enriched | Not Enriched |
| GO:0044718 | siderophore transmembrane transport | 2 | Not Enriched | Enriched | Not Enriched | Enriched | Not Enriched | Not Enriched |

| | | | | | | | | |
|---|---|---|---|---|---|---|---|---|
| GO:0006097 | glyoxylate cycle | 2 | Enriched | Not Enriched | Not Enriched | Enriched | Not Enriched | Not Enriched |
| GO:0097428 | protein maturation by iron-sulfur cluste... | 2 | Not Enriched | Not Enriched | Not Enriched | Enriched | Enriched | Not Enriched |
| GO:0065008 | regulation of biological quality | 2 | Not Enriched | Not Enriched | Not Enriched | Enriched | Enriched | Not Enriched |
| GO:0015949 | nucleobase-containing small molecule int... | 2 | Not Enriched | Not Enriched | Not Enriched | Enriched | Enriched | Not Enriched |
| GO:0006812 | monoatomic cation transport | 2 | Not Enriched | Enriched | Not Enriched | Enriched | Not Enriched | Not Enriched |
| GO:0015768 | maltose transport | 2 | Enriched | Not Enriched | Not Enriched | Enriched | Not Enriched | Not Enriched |
| GO:0042956 | maltodextrin transmembrane transport | 2 | Enriched | Enriched | Not Enriched | Not Enriched | Not Enriched | Not Enriched |
| GO:0009409 | response to cold | 2 | Enriched | Enriched | Not Enriched | Not Enriched | Not Enriched | Not Enriched |
| GO:0052803 | imidazole-containing compound metabolic ... | 2 | Enriched | Not Enriched | Not Enriched | Not Enriched | Not Enriched | Enriched |
| GO:0000105 | histidine biosynthetic process | 2 | Enriched | Not Enriched | Not Enriched | Not Enriched | Not Enriched | Enriched |
| GO:0042135 | neurotransmitter catabolic process | 2 | Enriched | Not Enriched | Not Enriched | Not Enriched | Not Enriched | Enriched |
| GO:0005978 | glycogen biosynthetic process | 2 | Enriched | Enriched | Not Enriched | Not Enriched | Not Enriched | Not Enriched |
| GO:0034637 | cellular carbohydrate biosynthetic proce... | 2 | Not Enriched | Enriched | Not Enriched | Not Enriched | Enriched | Not Enriched |
| GO:0006094 | gluconeogenesis | 2 | Not Enriched | Not Enriched | Not Enriched | Not Enriched | Enriched | Enriched |
| GO:1901271 | lipooligosaccharide biosynthetic process | 2 | Not Enriched | Not Enriched | Not Enriched | Not Enriched | Enriched | Enriched |

| | | | | | | | | |
|---|---|---|---|---|---|---|---|---|
| GO:0009247 | glycolipid biosynthetic process | 2 | Not Enriched | Not Enriched | Not Enriched | Not Enriched | Enriched | Enriched |
| GO:0016226 | iron-sulfur cluster assembly | 1 | Not Enriched | Not Enriched | Enriched | Not Enriched | Not Enriched | Not Enriched |
| GO:0006879 | intracellular iron ion homeostasis | 1 | Not Enriched | Not Enriched | Enriched | Not Enriched | Not Enriched | Not Enriched |
| GO:0009242 | colanic acid biosynthetic process | 1 | Not Enriched | Not Enriched | Enriched | Not Enriched | Not Enriched | Not Enriched |
| GO:0051674 | localization of cell | 1 | Not Enriched | Not Enriched | Enriched | Not Enriched | Not Enriched | Not Enriched |
| GO:0032508 | DNA duplex unwinding | 1 | Not Enriched | Not Enriched | Enriched | Not Enriched | Not Enriched | Not Enriched |
| GO:0035672 | oligopeptide transmembrane transport | 1 | Not Enriched | Not Enriched | Enriched | Not Enriched | Not Enriched | Not Enriched |
| GO:0051130 | positive regulation of cellular componen... | 1 | Not Enriched | Not Enriched | Enriched | Not Enriched | Not Enriched | Not Enriched |
| GO:0071973 | bacterial-type flagellum-dependent cell ... | 1 | Not Enriched | Not Enriched | Enriched | Not Enriched | Not Enriched | Not Enriched |
| GO:0006006 | glucose metabolic process | 1 | Not Enriched | Not Enriched | Enriched | Not Enriched | Not Enriched | Not Enriched |
| GO:1902047 | polyamine transmembrane transport | 1 | Not Enriched | Not Enriched | Enriched | Not Enriched | Not Enriched | Not Enriched |
| GO:0051606 | detection of stimulus | 1 | Not Enriched | Not Enriched | Enriched | Not Enriched | Not Enriched | Not Enriched |
| GO:0046496 | nicotinamide nucleotide metabolic proces... | 1 | Not Enriched | Not Enriched | Enriched | Not Enriched | Not Enriched | Not Enriched |
| GO:0009267 | cellular response to starvation | 1 | Not Enriched | Not Enriched | Enriched | Not Enriched | Not Enriched | Not Enriched |

| | | | | | | | | |
|---|---|---|---|---|---|---|---|---|
| GO:0051128 | regulation of cellular component organiz... | 1 | Not Enriched | Not Enriched | Enriched | Not Enriched | Not Enriched | Not Enriched |
| GO:0006113 | fermentation | 1 | Not Enriched | Not Enriched | Not Enriched | Enriched | Not Enriched | Not Enriched |
| GO:0019541 | propionate metabolic process | 1 | Not Enriched | Not Enriched | Not Enriched | Enriched | Not Enriched | Not Enriched |
| GO:0043436 | oxoacid metabolic process | 1 | Not Enriched | Not Enriched | Not Enriched | Enriched | Not Enriched | Not Enriched |
| GO:0022904 | respiratory electron transport chain | 1 | Not Enriched | Not Enriched | Not Enriched | Enriched | Not Enriched | Not Enriched |
| GO:0017000 | antibiotic biosynthetic process | 1 | Not Enriched | Not Enriched | Not Enriched | Enriched | Not Enriched | Not Enriched |
| GO:0033036 | macromolecule localization | 1 | Not Enriched | Not Enriched | Not Enriched | Enriched | Not Enriched | Not Enriched |
| GO:0015944 | formate oxidation | 1 | Not Enriched | Not Enriched | Not Enriched | Enriched | Not Enriched | Not Enriched |
| GO:2001057 | reactive nitrogen species metabolic proc... | 1 | Not Enriched | Not Enriched | Not Enriched | Enriched | Not Enriched | Not Enriched |
| GO:0007155 | cell adhesion | 1 | Not Enriched | Not Enriched | Not Enriched | Enriched | Not Enriched | Not Enriched |
| GO:0044780 | bacterial-type flagellum assembly | 1 | Not Enriched | Not Enriched | Not Enriched | Enriched | Not Enriched | Not Enriched |
| GO:0031669 | cellular response to nutrient levels | 1 | Not Enriched | Not Enriched | Not Enriched | Enriched | Not Enriched | Not Enriched |
| GO:0032465 | regulation of cytokinesis | 1 | Not Enriched | Not Enriched | Not Enriched | Enriched | Not Enriched | Not Enriched |
| GO:0009092 | homoserine metabolic process | 1 | Not Enriched | Not Enriched | Not Enriched | Enriched | Not Enriched | Not Enriched |
| GO:0072523 | purine-containing compound catabolic pro... | 1 | Not Enriched | Not Enriched | Not Enriched | Enriched | Not Enriched | Not Enriched |

| | | | | | | | | |
|---|---|---|---|---|---|---|---|---|
| GO:0006412 | translation | 1 | Enriched | Not Enriched | Not Enriched | Not Enriched | Not Enriched | Not Enriched |
| GO:0000028 | ribosomal small subunit assembly | 1 | Enriched | Not Enriched | Not Enriched | Not Enriched | Not Enriched | Not Enriched |
| GO:0000027 | ribosomal large subunit assembly | 1 | Enriched | Not Enriched | Not Enriched | Not Enriched | Not Enriched | Not Enriched |
| GO:0009097 | isoleucine biosynthetic process | 1 | Enriched | Not Enriched | Not Enriched | Not Enriched | Not Enriched | Not Enriched |
| GO:0009447 | putrescine catabolic process | 1 | Enriched | Not Enriched | Not Enriched | Not Enriched | Not Enriched | Not Enriched |
| GO:0015990 | electron transport coupled proton transp... | 1 | Enriched | Not Enriched | Not Enriched | Not Enriched | Not Enriched | Not Enriched |
| GO:0006637 | acyl-CoA metabolic process | 1 | Enriched | Not Enriched | Not Enriched | Not Enriched | Not Enriched | Not Enriched |
| GO:0009099 | valine biosynthetic process | 1 | Enriched | Not Enriched | Not Enriched | Not Enriched | Not Enriched | Not Enriched |
| GO:0008152 | metabolic process | 1 | Enriched | Not Enriched | Not Enriched | Not Enriched | Not Enriched | Not Enriched |
| GO:0046185 | aldehyde catabolic process | 1 | Enriched | Not Enriched | Not Enriched | Not Enriched | Not Enriched | Not Enriched |
| GO:1901607 | alpha-amino acid biosynthetic process | 1 | Enriched | Not Enriched | Not Enriched | Not Enriched | Not Enriched | Not Enriched |
| GO:0045947 | negative regulation of translational ini... | 1 | Enriched | Not Enriched | Not Enriched | Not Enriched | Not Enriched | Not Enriched |
| GO:0009098 | leucine biosynthetic process | 1 | Enriched | Not Enriched | Not Enriched | Not Enriched | Not Enriched | Not Enriched |
| GO:1990748 | cellular detoxification | 1 | Enriched | Not Enriched | Not Enriched | Not Enriched | Not Enriched | Not Enriched |
| GO:0044282 | small molecule catabolic process | 1 | Enriched | Not Enriched | Not Enriched | Not Enriched | Not Enriched | Not Enriched |

| | | | | | | | | |
|---|---|---|---|---|---|---|---|---|
| GO:0009636 | response to toxic substance | 1 | Enriched | Not Enriched | Not Enriched | Not Enriched | Not Enriched | Not Enriched |
| GO:0009450 | gamma-aminobutyric acid catabolic proces... | 1 | Enriched | Not Enriched | Not Enriched | Not Enriched | Not Enriched | Not Enriched |
| GO:0046436 | D-alanine metabolic process | 1 | Enriched | Not Enriched | Not Enriched | Not Enriched | Not Enriched | Not Enriched |
| GO:0034644 | cellular response to UV | 1 | Enriched | Not Enriched | Not Enriched | Not Enriched | Not Enriched | Not Enriched |
| GO:0097164 | ammonium ion metabolic process | 1 | Enriched | Not Enriched | Not Enriched | Not Enriched | Not Enriched | Not Enriched |
| GO:0034641 | cellular nitrogen compound metabolic pro... | 1 | Enriched | Not Enriched | Not Enriched | Not Enriched | Not Enriched | Not Enriched |
| GO:0017148 | negative regulation of translation | 1 | Enriched | Not Enriched | Not Enriched | Not Enriched | Not Enriched | Not Enriched |
| GO:0000162 | tryptophan biosynthetic process | 1 | Enriched | Not Enriched | Not Enriched | Not Enriched | Not Enriched | Not Enriched |
| GO:0009063 | amino acid catabolic process | 1 | Enriched | Not Enriched | Not Enriched | Not Enriched | Not Enriched | Not Enriched |
| GO:0050821 | protein stabilization | 1 | Not Enriched | Not Enriched | Not Enriched | Not Enriched | Enriched | Not Enriched |
| GO:0033214 | siderophore-dependent iron import into c... | 1 | Not Enriched | Not Enriched | Not Enriched | Not Enriched | Enriched | Not Enriched |
| GO:0051701 | biological process involved in interacti... | 1 | Not Enriched | Not Enriched | Not Enriched | Not Enriched | Enriched | Not Enriched |
| GO:0009432 | SOS response | 1 | Not Enriched | Not Enriched | Not Enriched | Not Enriched | Enriched | Not Enriched |
| GO:0009245 | lipid A biosynthetic process | 1 | Not Enriched | Not Enriched | Not Enriched | Not Enriched | Enriched | Not Enriched |
| GO:0046677 | response to antibiotic | 1 | Not Enriched | Not Enriched | Not Enriched | Not Enriched | Enriched | Not Enriched |

| | | | | | | | | |
|---|---|---|---|---|---|---|---|---|
| GO:0006073 | cellular glucan metabolic process | 1 | Not Enriched | Not Enriched | Not Enriched | Not Enriched | Enriched | Not Enriched |
| GO:0010468 | regulation of gene expression | 1 | Not Enriched | Not Enriched | Not Enriched | Not Enriched | Enriched | Not Enriched |
| GO:0007165 | signal transduction | 1 | Not Enriched | Not Enriched | Not Enriched | Not Enriched | Enriched | Not Enriched |
| GO:0006207 | de novo' pyrimidine nucleobase biosynth... | 1 | Not Enriched | Not Enriched | Not Enriched | Not Enriched | Enriched | Not Enriched |
| GO:0006083 | acetate metabolic process | 1 | Not Enriched | Not Enriched | Not Enriched | Not Enriched | Enriched | Not Enriched |
| GO:0010041 | response to iron(III) ion | 1 | Not Enriched | Not Enriched | Not Enriched | Not Enriched | Enriched | Not Enriched |
| GO:0046349 | amino sugar biosynthetic process | 1 | Not Enriched | Not Enriched | Not Enriched | Not Enriched | Enriched | Not Enriched |
| GO:0061077 | chaperone-mediated protein folding | 1 | Not Enriched | Not Enriched | Not Enriched | Not Enriched | Enriched | Not Enriched |
| GO:0051253 | negative regulation of RNA metabolic pro... | 1 | Not Enriched | Not Enriched | Not Enriched | Not Enriched | Not Enriched | Enriched |
| GO:0042866 | pyruvate biosynthetic process | 1 | Not Enriched | Not Enriched | Not Enriched | Not Enriched | Not Enriched | Enriched |
| GO:0006096 | glycolytic process | 1 | Not Enriched | Not Enriched | Not Enriched | Not Enriched | Not Enriched | Enriched |
| GO:0006754 | ATP biosynthetic process | 1 | Not Enriched | Not Enriched | Not Enriched | Not Enriched | Not Enriched | Enriched |
| GO:0019359 | nicotinamide nucleotide biosynthetic pro... | 1 | Not Enriched | Not Enriched | Not Enriched | Not Enriched | Not Enriched | Enriched |
| GO:0006090 | pyruvate metabolic process | 1 | Not Enriched | Not Enriched | Not Enriched | Not Enriched | Not Enriched | Enriched |
| GO:0009071 | serine family amino acid catabolic proce... | 1 | Not Enriched | Not Enriched | Not Enriched | Not Enriched | Not Enriched | Enriched |

| | | | | | | | | |
|---|---|---|---|---|---|---|---|---|
| GO:0042594 | response to starvation | 1 | Not Enriched | Not Enriched | Not Enriched | Not Enriched | Not Enriched | Enriched |
| GO:0009250 | glucan biosynthetic process | 1 | Not Enriched | Not Enriched | Not Enriched | Not Enriched | Not Enriched | Enriched |
| GO:0002098 | tRNA wobble uridine modification | 1 | Not Enriched | Not Enriched | Not Enriched | Not Enriched | Not Enriched | Enriched |
| GO:0006415 | translational termination | 1 | Not Enriched | Not Enriched | Not Enriched | Not Enriched | Not Enriched | Enriched |
| GO:0009166 | nucleotide catabolic process | 1 | Not Enriched | Not Enriched | Not Enriched | Not Enriched | Not Enriched | Enriched |
| GO:0034219 | carbohydrate transmembrane transport | 1 | Not Enriched | Enriched | Not Enriched | Not Enriched | Not Enriched | Not Enriched |
| GO:0031460 | glycine betaine transport | 1 | Not Enriched | Enriched | Not Enriched | Not Enriched | Not Enriched | Not Enriched |
| GO:0006820 | monoatomic anion transport | 1 | Not Enriched | Enriched | Not Enriched | Not Enriched | Not Enriched | Not Enriched |
| GO:0009401 | phosphoenolpyruvate-dependent sugar phos... | 1 | Not Enriched | Enriched | Not Enriched | Not Enriched | Not Enriched | Not Enriched |
| GO:0071214 | cellular response to abiotic stimulus | 1 | Not Enriched | Enriched | Not Enriched | Not Enriched | Not Enriched | Not Enriched |
| GO:0006836 | neurotransmitter transport | 1 | Not Enriched | Enriched | Not Enriched | Not Enriched | Not Enriched | Not Enriched |
| GO:0097305 | response to alcohol | 1 | Not Enriched | Enriched | Not Enriched | Not Enriched | Not Enriched | Not Enriched |
| GO:0006596 | polyamine biosynthetic process | 1 | Not Enriched | Enriched | Not Enriched | Not Enriched | Not Enriched | Not Enriched |
| GO:0015757 | galactose transmembrane transport | 1 | Not Enriched | Enriched | Not Enriched | Not Enriched | Not Enriched | Not Enriched |

| | | | | | | | | |
|---|---|---|---|---|---|---|---|---|
| GO:0051085 | chaperone cofactor-dependent protein ref... | 1 | Not Enriched | Enriched | Not Enriched | Not Enriched | Not Enriched | Not Enriched |
| GO:0019579 | aldaric acid catabolic process | 1 | Not Enriched | Enriched | Not Enriched | Not Enriched | Not Enriched | Not Enriched |
| GO:0019220 | regulation of phosphate metabolic proces... | 1 | Not Enriched | Enriched | Not Enriched | Not Enriched | Not Enriched | Not Enriched |
| GO:0046031 | ADP metabolic process | 1 | Not Enriched | Enriched | Not Enriched | Not Enriched | Not Enriched | Not Enriched |
| GO:0032879 | regulation of localization | 1 | Not Enriched | Enriched | Not Enriched | Not Enriched | Not Enriched | Not Enriched |
| GO:0033554 | cellular response to stress | 1 | Not Enriched | Enriched | Not Enriched | Not Enriched | Not Enriched | Not Enriched |
| GO:0097237 | cellular response to toxic substance | 1 | Not Enriched | Enriched | Not Enriched | Not Enriched | Not Enriched | Not Enriched |
| GO:0033499 | galactose catabolic process via UDP-gala... | 1 | Not Enriched | Enriched | Not Enriched | Not Enriched | Not Enriched | Not Enriched |
| GO:0015766 | disaccharide transport | 1 | Not Enriched | Enriched | Not Enriched | Not Enriched | Not Enriched | Not Enriched |
| GO:0042940 | D-amino acid transport | 1 | Not Enriched | Enriched | Not Enriched | Not Enriched | Not Enriched | Not Enriched |
| GO:0032101 | regulation of response to external stimu... | 1 | Not Enriched | Enriched | Not Enriched | Not Enriched | Not Enriched | Not Enriched |
| GO:1902021 | regulation of bacterial-type flagellum-d... | 1 | Not Enriched | Enriched | Not Enriched | Not Enriched | Not Enriched | Not Enriched |
| GO:0070887 | cellular response to chemical stimulus | 1 | Not Enriched | Enriched | Not Enriched | Not Enriched | Not Enriched | Not Enriched |
| GO:0006089 | lactate metabolic process | 1 | Not Enriched | Enriched | Not Enriched | Not Enriched | Not Enriched | Not Enriched |
| GO:0044403 | biological process involved in symbiotic... | 1 | Not Enriched | Enriched | Not Enriched | Not Enriched | Not Enriched | Not Enriched |

| | | | | | | | | |
|---|---|---|---|---|---|---|---|---|
| GO:0032774 | RNA biosynthetic process | 1 | Not Enriched | Enriched | Not Enriched | Not Enriched | Not Enriched | Not Enriched |
| GO:0046835 | carbohydrate phosphorylation | 1 | Not Enriched | Enriched | Not Enriched | Not Enriched | Not Enriched | Not Enriched |
| GO:0044262 | cellular carbohydrate metabolic process | 1 | Not Enriched | Enriched | Not Enriched | Not Enriched | Not Enriched | Not Enriched |
| GO:0008645 | hexose transmembrane transport | 1 | Not Enriched | Enriched | Not Enriched | Not Enriched | Not Enriched | Not Enriched |
| GO:0070301 | cellular response to hydrogen peroxide | 1 | Not Enriched | Enriched | Not Enriched | Not Enriched | Not Enriched | Not Enriched |

**Table S4:** Strains used in this study

| Species | Published strain number | Lab isolate number (if different) | Genotype | Source | Reference | NCBI assembly used for RNA-seq analysis |
|---|---|---|---|---|---|---|
| *E. coli* | BW27786 | DMS2537 | F-, Δ(araD-araB)567, ΔlacZ4787(::rrnB-3), λ-, Δ(araH-araF)570(::FRT), ΔaraEp-532::FRT, φPcp13araE534, Δ(rhaD-rhaB)568, hsdR514 | Lab collection | doi: 10.1099/00221287-147-12-3241 | GCF_000005845.2 |
| *E. coli* | DMS2545 | | ΔrpoS746::kan derivative of DMS2537 (F-, Δ(araD-araB)567, ΔlacZ4787(::rrnB-3), λ-, Δ(araH-araF)570(::FRT), ΔaraEp-532::FRT, φPcp13araE534, Δ(rhaD-rhaB)568, hsdR514, ΔrpoS746::kan) | Lab collection | doi: 10.1128/jb.00755-16 | GCF_000005845.2 |
| *S. enterica* | 14028s | DMS2709 | wild-type | Salmonella Genetic Stock Center | | GCF_000022165.1 |
| *S. enterica* | | DMS3189 | Δ*rpoS*::cm derivative of 14028s | McClelland Lab | | GCF_000022165.1 |
| *C. rodentium* | DBS100 | DMS2868 | wild-type | ATCC | | GCF_000027085.1 |
| *E. cloacae* | ATCC13047 | DMS2862 | wild-type | ATCC | | GCF_000025565.1 |
| *K. pneumoniae* | MKP103 | DMS2797 | Strain KPNIH01, Δ*KPC-3::FRT* | Manoil Lab | doi: 10.1128/JB.00352-17 | GCF_000281535.2 |
| *S. marcescens* | Db11 | DMS2843 | wild-type | *C. elegans* Stock Center | | GCF_000513215.1 |

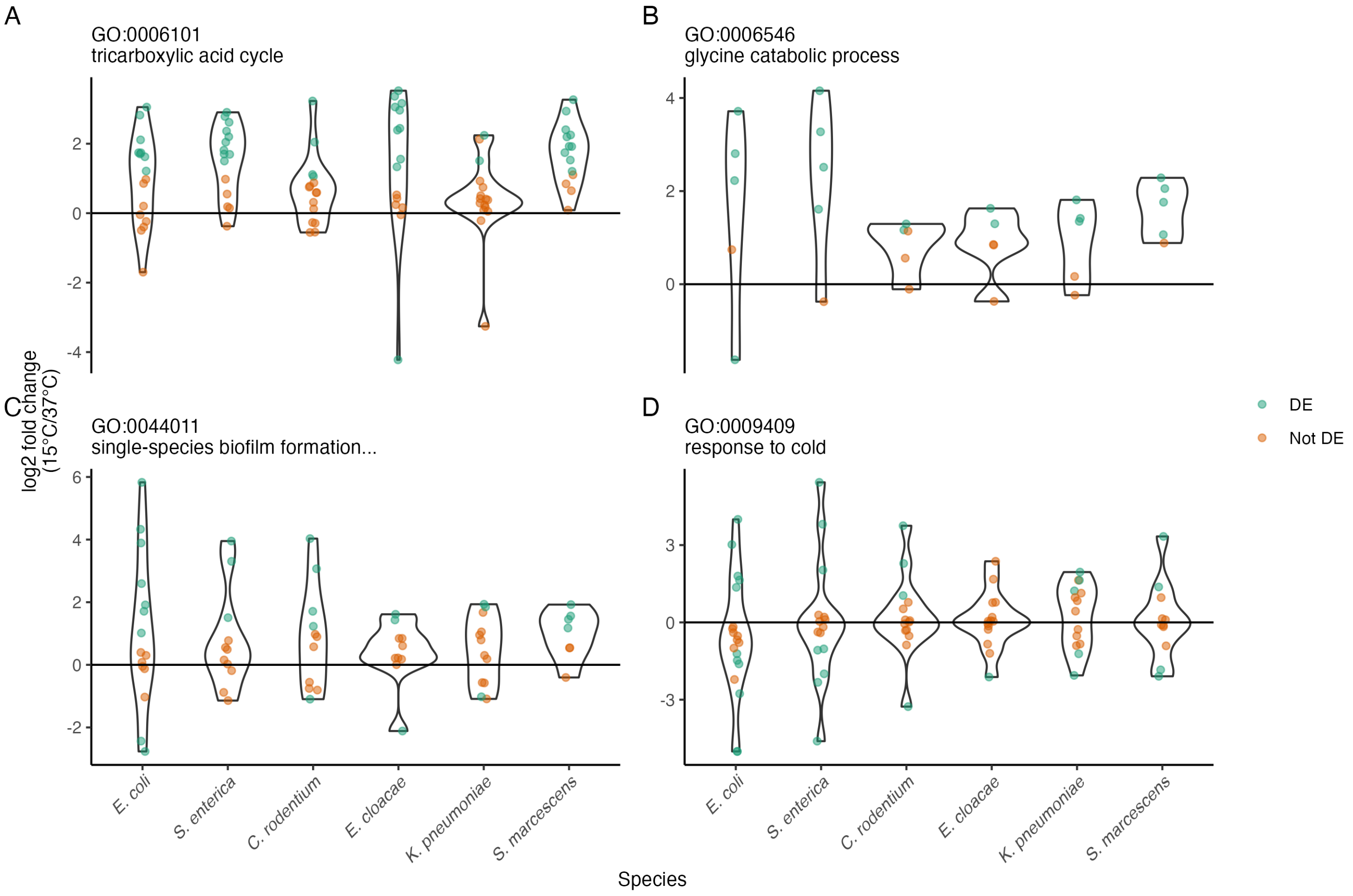


**Figure S1: Groups of genes that change in multiple species.** $Log_2$-fold change of gene expression between 15°C and 37°C for genes annotated as involved in (A) the tricarboxylic acid cycle, (B) glycine catabolism, (C) single-species biofilm formation on an inanimate surface, and (D) response to cold. Green dots represent a single DE gene, while orange dots are a gene that is not DE. Violin plots show the distributions of $log_2$-fold changes of the individual genes.

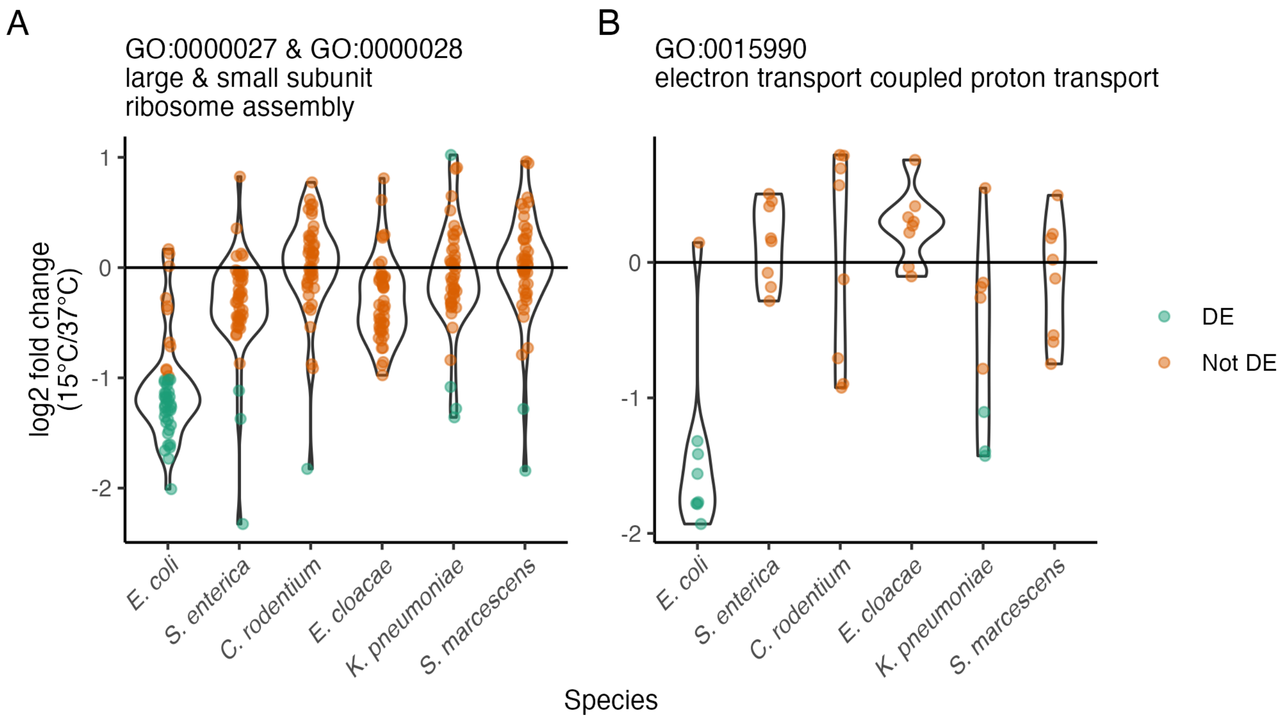


**Figure S2: Responses to low temperature unique to *E. coli*.** Log$_2$-fold change of gene expression between 15°C and 37°C for genes annotated as involved in (A) production of the ribosome, and (B) electron transport coupled proton transport. Green dots represent a single DE gene, while orange dots are a gene that is not DE. Violin plots show the distributions of log$_2$-fold changes of the individual genes.

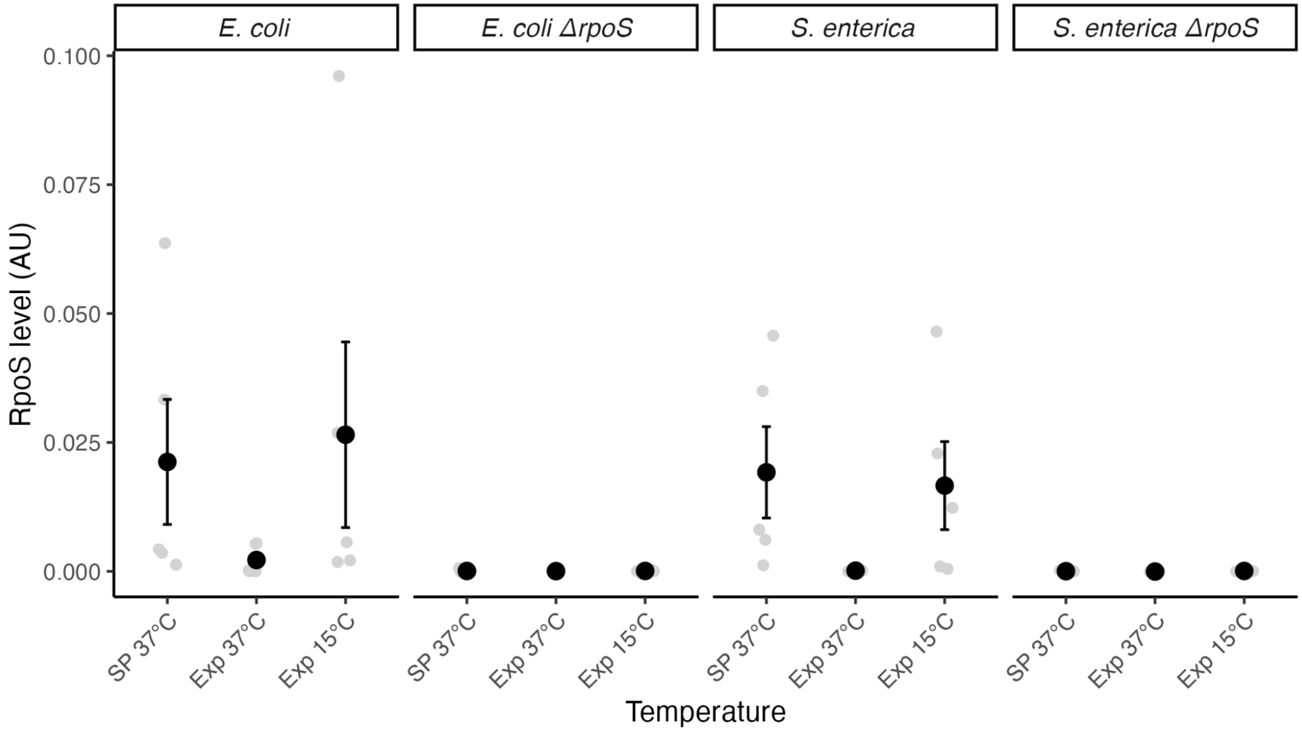


**Figure S3:** RpoS levels as measured by Western blotting in stationary phase (SP), exponentially growing cultures at 37°C (Exp 37°C), or of those same cultures 3 hours after a shift to 15°C (Exp 15°C). Light gray dots are individual replicates, black dots are the mean, and error bars are the standard error of the mean.
